# HIF-1α and RhoA Drive Enhanced Motility and Aerotaxis of Polyaneuploid Prostate Cancer Cells in Hypoxia

**DOI:** 10.1101/2025.09.03.673887

**Authors:** Noreen Hosny, Shengkai Li, Sarah Amend, Robert Gatenby, Kenneth J. Pienta, Joel Brown, Junle Qu, Robert Austin

## Abstract

Most cancer deaths result from metastasis, yet only a rare subset of tumor cells can complete this process. Among these, polyaneuploid cancer cells (PACCs), which arise via endoreplication under stressors such as hypoxia, are implicated as metastatic drivers, but how they acquire this potential is poorly understood. Here, we show that prostate cancer-derived PACCs exhibit features predictive of invasion and intravasation. Time-lapse fluorescence microscopy and single-cell tracking under hypoxia revealed that PACCs migrated significantly farther than Non-PACCs, consistent with local invasion. PACC trajectories showed a strong tendency to migrate toward oxygen, consistent with aerotaxis and predictive of intravasation. siRNA-mediated knockdown demonstrated that the enhanced motility and aerotaxis of hypoxic PACCs require both HIF-1αand RhoA, with RhoA expression suppressed upon HIF-1α inhibition. Altogether, we propose a HIF-1α→ RhoA → motility/aerotaxis mechanism enabling PACCs to escape hypoxic cores, invade tissue, and access vasculature, highlighting them as a uniquely invasive subpopulation with implications for anti-metastatic therapies.

## INTRODUCTION

Metastasis describes the development of secondary malignant tumors at a distant site from the primary cancer, and is responsible for over 90% of cancer-related deaths.^1^ The multistep, metastatic cascade begins with the local invasion of cancer cells into adjacent tissues, a key step often marked by the epithelial to mesenchymal transition (EMT). ^2,3^ Cancer cells then proceed through the subsequent phases of the metastatic cascade: intravasation into nearby blood vessels, entrance into the circulatory system as circulating tumor cells (CTCs), survival within the bloodstream, extravasation into new tissues as disseminating tumor cells (DTCs), and finally, colonization at secondary sites to form new malignant growths. ^2–6^

Despite metastasis being the leading cause of death in cancer patients, only a select few cancer cells are capable of completing the full metastatic cascade to form clinically-detectable lesions. Preclinical studies in melanoma, breast, and prostate cancer models estimate that while approximately one million tumor cells per gram of primary tumor enter circulation each day, fewer than 0.01% ultimately give rise to metastases.^7^ This inefficiency reflects the immense physiological barriers imposed by the cascade; only a small fraction of these cells can withstand the shear stress of blood flow and adapt to the distinct microenvironments of secondary sites.^8^ Thus, characterization of the cancer cells with uniquely high metastatic potential is crucial for elucidating the mechanisms that drive metastasis at the single-cell level.

Recent research has suggested that polyaneuploid cancer cells (PACCs), also termed polyploid giant cancer cells (PGCCs), possess this increased capacity for metastasis.^9,10^ Once believed to be benign and permanently senescent, PACCs are now recognized as highly plastic, stress-adapted cells that contribute to cancer lethality.^9,11–14^ The PACC state arises when an aneuploid parent cell undergoes endoreplication in response to environmental stressors such as hypoxia, chemotherapy, or nutrient deprivation, resulting in extensive polyploidization.^9,15,16^ The resulting genomic amplification is thought to enhance the biosynthetic and DNA repair capacities of PACCs, promoting the evasion of mitotic catastrophe and enabling persistence under adverse conditions.^17^

Clinically, cells in the PACC state have been identified in the malignant solid tumors of several cancer types, including melanoma, pancreatic, ovarian, kidney, and prostate cancer.^11–14^ Further alluding to the potential metastatic nature of PACCs, these cells are more common in secondary metastatic sites rather than the primary tumor.^11,12^ Preliminary *in-vitro* studies using cisplatin-treated PC3 prostate cancer cells that entered the PACC state exhibited increased motility, thereby suggesting an increased ability to successfully complete invasion during metastasis.^9,10^ Despite this growing evidence that PACCs contribute to metastatic progression, the physical characteristics and molecular mechanisms that confer this metastatic potential remain poorly understood.

To address this, we sought to characterize PACCs that emerge specifically in response to hypoxia, defined as oxygen concentrations falling below ∼2%.^18^

Hypoxia, a hallmark of solid tumors, imposes strong selective pressures on cancer cells and has been linked to increased motility and invasiveness.^19–21^ Beyond metabolic reprogramming, hypoxia may stimulate aerotaxis, the directed migration of cells toward oxygen-rich regions, potentially facilitating intravasation.^22^ Whether cancer cells in the PACC state exhibit increased aerotactic migration remains another open question.

Hypoxia-inducible factors (HIFs) play a central role in cancer cell adaptation to low oxygen and in promoting invasiveness. Under normoxia, HIF-1αis targeted for proteasomal degradation by prolyl hydroxylases, but in hypoxia it becomes stabilized, translocates to the nucleus, and dimerizes with HIF-1β.^23^ This complex then activates genes that express glycolytic enzymes, facilitating the metabolic shift toward anaerobic glycolysis.^23,24^ HIF-1αalso upregulates genes that drive epithelial-to-mesenchymal transition (EMT) and thus promote invasiveness.^20,21^ Similarly, RhoA, a Rho GTPase, promotes cell motility by remodeling the actin cytoskeleton through activation of ROCK1/2, driving actomyosin contractility.^25^ Notably, both RhoA and ROCK kinases are frequently upregulated in cancers such as renal, breast, and colon.^26–28^ Thus, HIF-1αand RhoA represent complementary mechanisms for promoting cancer cell motility, which may also support PACC behavior under hypoxia. Given the overlapping roles of HIF-1αand RhoA, we sought to investigate whether a mechanistic connection exists between them. Prior studies suggest context-dependent directionality, with HIF-1αacting upstream of RhoA in colon cancer but the reverse in renal cell carcinoma, raising the possibility of cell type-specific variability.^29,30^

Here, we show that PACC motility and aerotaxis are enhanced under hypoxia via live cell microscopy and single-cell tracking. siRNA knockdown and fixed-cell fluorescence microscopy revealed that HIF-1αand RhoA are required for these behaviors, and that RhoA expression is HIF-1α-dependent in hypoxic prostate cancer cells. Together, these findings support a model in which HIF-1αactivation of RhoA drives PACC motility and oxygen-directed migration, enabling these cells to invade surrounding tissue and access the oxygen-rich vasculature. Collectively, our results identify PACCs as a highly invasive subpopulation and highlight the HIF-1α-RhoA pathway as a potential target for limiting metastatic spread in prostate cancer.

## RESULTS

### Hypoxia promotes the emergence of PACCs

PACCs have been characterized as large, stress-resistant cancer cells that arise predominantly through endoreplication, leading to increased genomic content and cell size.^9,14–17,31^ To assess whether hypoxia promotes the PACC state and to evaluate their motility and molecular characteristics under physiologically relevant conditions, we used the membrane-based culture system employed in our previous study.^18^ This platform establishes a stable oxygen gradient across the culture dish and allows spatial visualization of hypoxia development.^18^ Using this system, we examined whether hypoxia induces the PACC state through immunofluorescence imaging of cell and nuclear size, together with FACS analysis of DNA content following 16 hours of hypoxia. Consistent with previous reports, we observed cells with markedly enlarged nuclei and overall increased cell size among a population of smaller cells in the hypoxia condition, in contrast to normoxic cells, which displayed relatively uniform and smaller nuclear and cytoplasmic areas (Fig. 1A-F).^14,31–33^ FACS analysis revealed a nearly threefold increase in the proportion of PACCs under hypoxic conditions compared to normoxia, based on identification of cells with greater than 4N DNA content (Fig. 1G-I). Corresponding cell cycle distributions further demonstrated a rightward shift beyond the G2/M peak (Supplementary Fig. S1). Taken together, these results suggest that the membrane-based culture system recapitulates tumor heterogeneity and promotes the hypoxia-driven enrichment of the PACC population.

**Figure 1.**
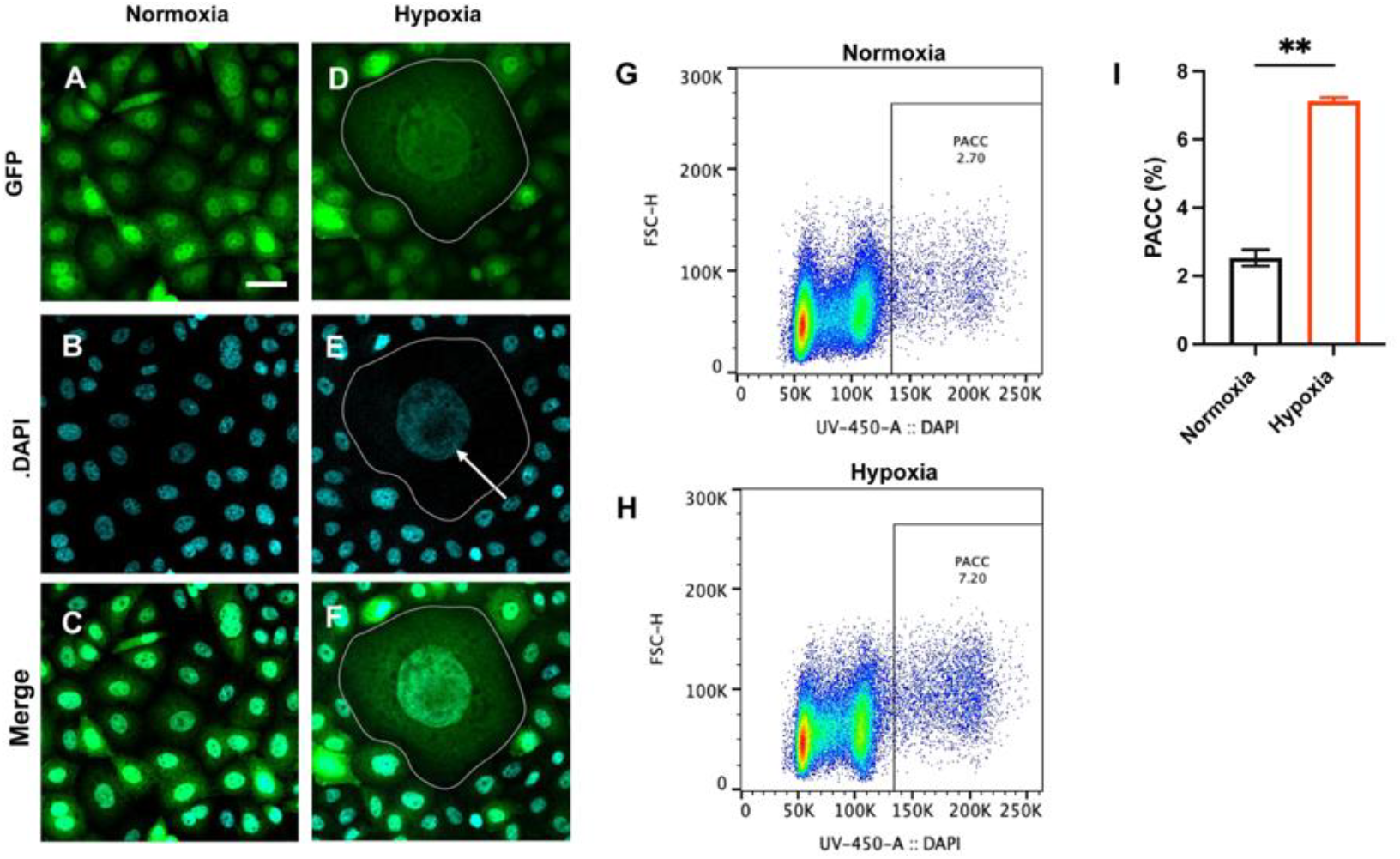
Hypoxia promotes the emergence of PACCs. **(A-C)** Representative image of PC3-GFP cells under normoxic conditions. **(D-F)** Representative image of PC3-GFP cells after 16 hours of hypoxia, with PACC cell (outlined) surrounded by non-PACC cells. The enlarged nucleus of the PACC cell is indicated by an arrow in **(E). (G, H)** Fluorescence-Activated Cell Sorting (FACS) analysis of PC3-GFP cells in normoxia **(G)** and hypoxia **(H).** PACCs were defined as cells with DNA content >4N. FSC-H: forward scatter height (proxy of cell size); UV- 450-A, DAPI fluorescence intensity (proxy for genomic content). **(I)** Quantification of PACCs from FACS analysis in normoxic and hypoxic conditions. Data are shown as mean ± SD. Significance determined in **(I)** as p ≤ 0.01 (^**^) via Unpaired Student’s t test (see Material and Methods). Scale bars for **A-F** are in **A** and represent 20 μm.

### PACCs exhibit increased motility in hypoxia

Motility is essential for cancer cells to penetrate the basement membrane that separates the primary tumor from surrounding tissues.^2–4,8^ Once this barrier is breached, cancer cells can initiate the metastatic cascade by moving into blood vessels (intravasation) and traveling to distant sites.^2,8,34^ Thus, to assess the motility of PC3-GFP cells within self-generated O_2_ gradients, we performed time-lapse fluorescence microscopy followed by single-cell tracking using the TrackMate and Cellpose plugins in ImageJ (see Methods) to derive cell trajectories and cell areas over 16 hours of hypoxia (Figure 2A, B). Cells selected for analysis were located within regions of the steepest O_2_ gradient.

**Figure 2.**
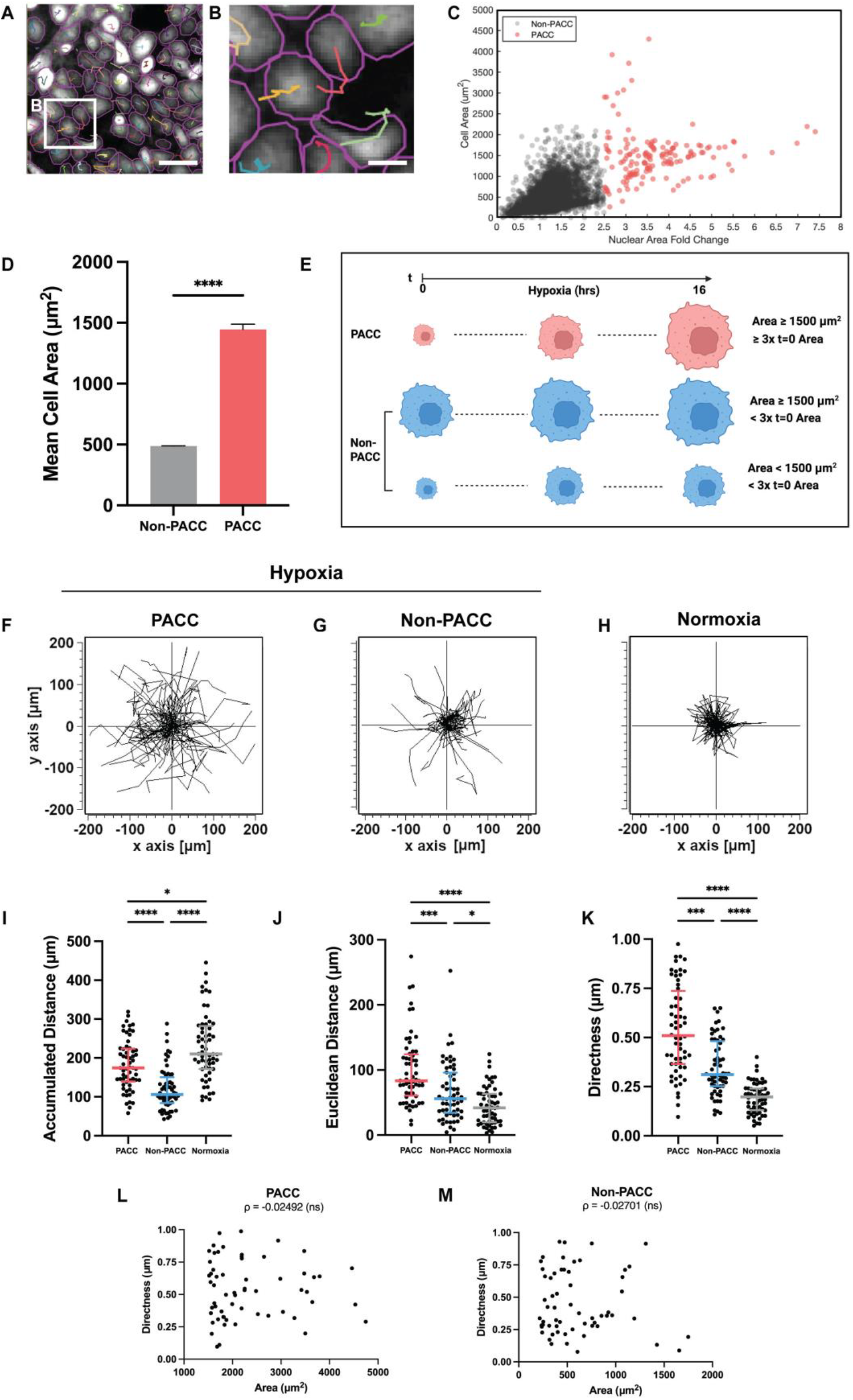
PACCs exhibit increased motility in hypoxia. (**A, B)** Detection of cell trajectories (colored tracks) using Trackmate prior to trajectory analysis. Cell segmentation (magenta) was conducted using Cellpose detector module within Trackmate. **(C)** Single-cell quantification of nuclear area fold change (relative to the mean nuclear area of the parent cell population, 116.55 μm^2^) versus cell area (µm^2^) of Non-PACCs (red) and PACCs (black). PACCs were defined as cells exceeding a nuclear area fold change of ≥2.5. Non-PACCs were defined as cells with a nuclear area fold change of < 2.5. Data pooled from three independent experiments. **(D)** Quantification of mean Non-PACC and PACC cell areas depicted in **(C).** Mean cell area of Non-PACC population, 487 μm^2^ ± 3 µm^2^, and that of PACC population is 1490 ± 48 μm^2^. Data are shown as Mean ± SEM. **(E)** Criteria for classifying PACC and Non-PACC cells in the hypoxic condition of single-cell tracking experiments. Cells that had an area of ≥1500 μm^2^ at t = 16 hours of hypoxia and grew at ≥3 times its cell area at t = 0 hours were defined as PACCs. Cells that had an area of ≥1500 μm^2^ at t = 16 hours of hypoxia, but grew < 3 times its initial cell area were defined as Non-PACCs. Cells that had an area of <1500 μm^2^ at t = 16 hours of hypoxia, and grew < 3 times its initial cell area were also defined as Non-PACCs. **(F-H)** Spider plots depicting the trajectories of n = 60 PACC **(F)** and Non-PACC **(G)** in hypoxia, and control PC3-GFP cells in Normoxia **(H)** over 16 hours of hypoxia. n= 20 cells were sampled for each of 3 independent experiments. **(I-K)** Single-cell quantification of cell trajectories. Accumulated Distance **(I)**, Euclidean Distance **(J)**, and Directness **(K)** of n = 60, PACC and Non-PACC cells under hypoxia, and n = 60 control cells in Normoxia. n= 20 cells were sampled per group from each of 3 independent experiments. Data are shown as Median ± IQR. **(L**,**M)** Scatterplots of Directness vs Cell Area and Spearman’s ρ for PACC **(L)** and Non-PACC **(M)** cells. Significance determined in **(D)** via Welch’s t test, and in **(I-K)** via Kruskal-Wallis test with Dunn’s post hoc test; p < 0.05 (*), p <0.01 (^**^), p <0.001 (^***^), p <0.0001 (^****^), ns = not significant. Scale bar for **A**, 50 µm and for **B**, 20 µm.

While PACCs are traditionally identified by their increased nuclear area (>2.5x that of the parent population), we omitted nuclear staining during time-lapse experiments to avoid potential disruption of DNA replication, as reported in the literature.^31–33,35–37^ Thus, we sought to establish a cell area-based criterion for PACC identification. After 16 hours of hypoxia, cells were fixed, stained with DAPI, and then analyzed using Cellpose/TrackMate (see Methods) to quantify nuclear and cell areas. Figure 2C depicts the nuclear area fold change (relative to the normoxic parent population) versus total cell area. Supplementary Figure S2 shows the nuclear versus cell area scatterplot used to derive parent population measurements. When the well-established cutoff of 2.5x nuclear area was applied, two relatively distinct clusters emerged, with cells exceeding the nuclear size threshold exhibiting an average cell area of approximately 1500 μm^2^ (Fig. 2C, D). Based on these results, cells with a final area ≥1500 μm^2^ at 16 hours were classified as PACCs in the time-lapse experiments. However, given sample heterogeneity, we recognized that some large cells may stochastically reach this area threshold without undergoing true PACC formation. Previous reports have noted that PACCs exhibit rapid growth under hypoxic or chemotherapeutic stress, expanding to approximately three times their initial size.^14,31,33^ Therefore, we incorporated an additional dynamic criterion: cells were classified as PACCs if they reached ≥1500 μm^2^ *and* exhibited at least a threefold increase in area over the 16-hour period (Fig. 2E). Cells that reached ≥1500 μm^2^ but exhibited less than a threefold area expansion, as well as those with final areas <1500 μm^2^ and less than threefold growth, were classified as “Non-PACCs” (Fig. 2E).

Multiple parameters were then used to characterize motility across PACC and Non-PACC populations. Accumulated Distance (total distance traveled) was measured to capture general motility. To assess more direct and penetrative behavior, we quantified Euclidean Distance (net displacement) and Directness (Euclidean Distance divided by Accumulated Distance), where a Directness value of 1 indicates perfectly straight trajectories and 0 indicates highly circuitous paths.

Spider plots mapping cell trajectories with t=0 coordinates fixed at the origin (0,0) revealed that PACCs exhibited the longest and most direct movements, while Non-PACC cells showed shorter and less directed trajectories (Fig. 2F,G). Indeed, PACCs exhibited significantly greater Accumulated Distance, Euclidean Distance, and Directness compared to other groups under hypoxia (Fig. 2I-K). Interestingly, when compared to normoxic controls, cells exhibited higher Accumulated Distances under normoxia, likely due to the availability of O_2_ supporting maximal metabolic activity (Fig. 2H,I). However, Euclidean Distance and Directness remained significantly lower under this control condition relative to hypoxic PACC and Non-PACC cells, suggesting that hypoxia promotes more directed motility and is the most enhanced in PACCs (Fig. 2J, K).

To determine whether the enhanced motility of PACCs was merely a function of their large size, we performed correlational analyses between cell area and motility metrics using Spearman’s rank correlation coefficient (ρ) (Fig. 2L,M, Supplementary Fig. S3). Across all parameters, non-significant ρ coefficients indicated no correlation between cell size and Accumulated Distance, Euclidean Distance, or Directness in either PACCs or Non-PACCs (Fig. 2L,M, Supplementary Fig. S3), suggesting that the observed motility differences are largely independent of cell size and may instead be attributed to distinct molecular mechanisms under hypoxia.

### PACC cells exhibit increased aerotactic migration in hypoxia

In addition to general motility, the ability of cancer cells to undergo aerotactic migration, or directed movement toward O_2_-rich regions, may contribute to increasing metastatic potential.^2,22^ After breaching the basement membrane, cancer cells must continue migrating toward oxygenated blood vessels to complete intravasation.^2,3,34^ Recent research suggests that aerotactic migration can facilitate this process and also serve as a survival mechanism, allowing cells to escape the hypoxic tumor core.^22^

Thus, we sought to assess whether PACCs have an enhanced ability to sense the self-generated O_2_ gradient and migrate towards O_2_-rich regions. As a departure from traditional chemotaxis experiments that involve a uniform linear gradient across the entirety of the dish, our culture system generates a radial O_2_ gradient.^9,38^ As a result, each cell experiences a distinct local gradient that depends on its position in the dish (Fig. 3A). To enable direct comparison of migration behavior across cells, we rotated each cell’s trajectory such that its local O_2_ gradient was aligned with the positive y-direction (Fig. 3A, see Methods). This transformation allowed us to place all cells into a common reference frame, making it possible to assess whether movement was generally oriented toward O_2_-rich regions (Fig. 3A, see Methods).

**Figure 3.**
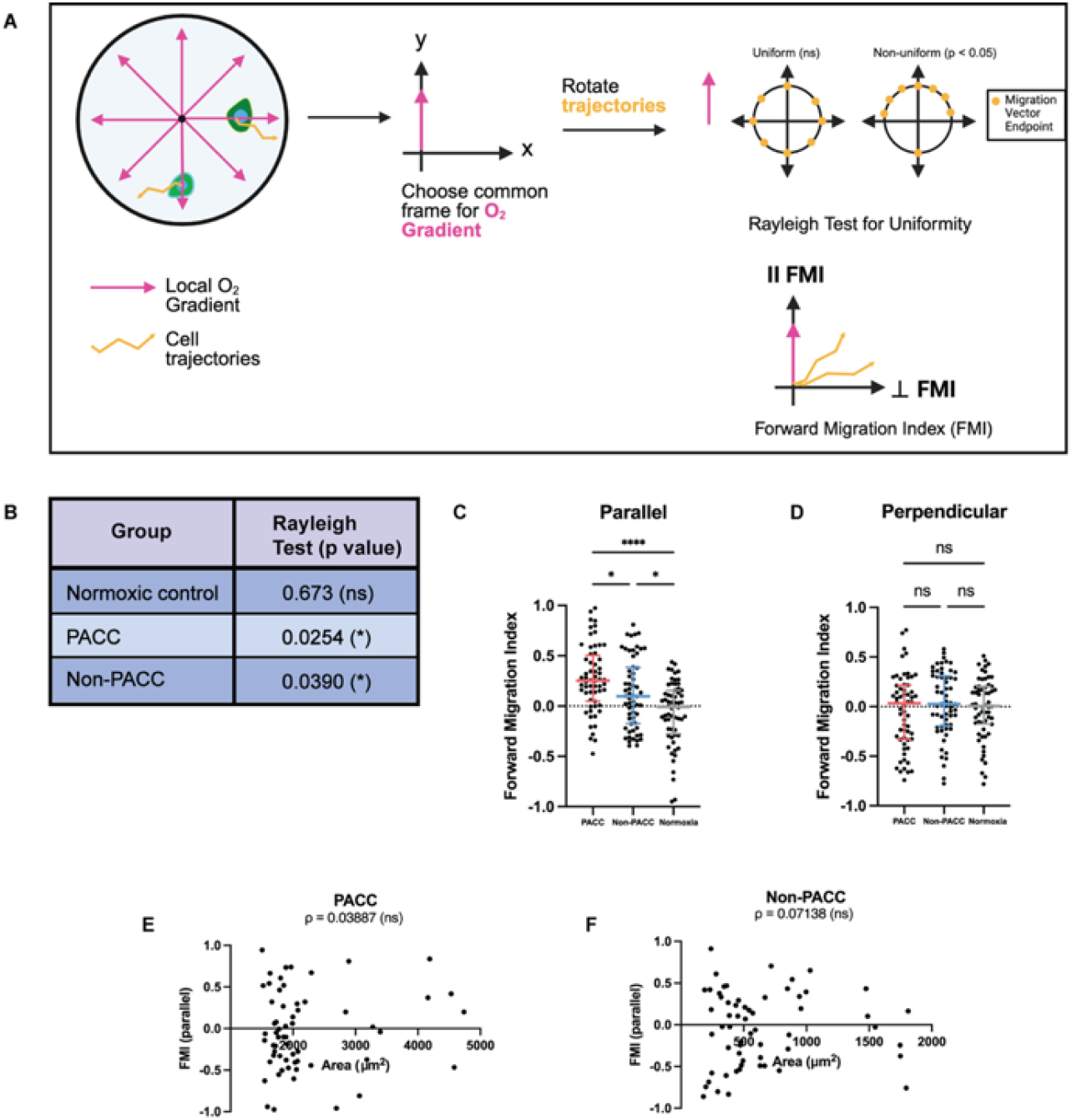
PACC cells exhibit enhanced aerotactic migration in hypoxia. **(A)** Schematic illustrating how cell trajectories were transformed into a common reference frame for quantifying directional migration. In our membrane-based culture system, the self-generated O_2_ gradient is radial (magenta arrows), meaning that each cell experiences a unique local gradient depending on its position. To standardize analyses, trajectories were rotated such that each local gradient vector was aligned with the positive y-axis. This transformation enabled evaluation of directional bias using the Rayleigh test for uniformity, as well as quantification of migration parallel (‖ Forward Migration Index, FMI) and perpendicular to (⊥ FMI) the O_2_ gradient (see Methods). **(B)** Rayleigh test p-values for each condition: PACC and Non-PACC Cells (under hypoxia), and Normoxic controls. A significant p-value indicates non-uniform (i.e., directional) migration. **(C, D)** ‖ FMI **(C)** and ⊥ FMI **(D)** relative to the O_2_ gradient across groups. The ‖ FMI reflects movement toward or away from the O_2_ source, while the ⊥ FMI captures lateral (non-directional) migration. Data are shown as Median ± IQR. **(E**,**F)** Scatterplots of ‖ FMI vs Cell Area and Spearman’s ρ for PACC **(E)** and Non-PACC **(F)** Cells. Data in **(B-F)** were acquired from n = 60 cells per group. Significance determined as p < 0.05 (*), p < 0.01 (^**^), p < 0.001 (^***^), and p < 0.0001 (^****^), and ns = not significant via Rayleigh Test in **(B)**, Kruskal-Wallis test followed by Dunn’s post hoc test in **(C, D)** (see Material and Methods).

To determine whether the cells exhibited O_2_-directed migration of statistical significance, we then computed the Rayleigh test for uniformity under the modified reference frame, in which all local O_2_ gradients were aligned along the positive y-axis. The Rayleigh test evaluates whether directional data are randomly distributed around a circle.^39,40^ In this context, the test was applied to the final displacement angles derived from spider plots, treating each trajectory as a vector originating from the origin. A nonsignificant result indicates a uniform, non-directional distribution of movement, whereas a significant result reflects a biased, directional response. Figure 3B displays the resulting p values of the test, which revealed significant directional migration in hypoxic PACC (p = 0.0254) and Non-PACC (p = 0.0390) cells, whereas normoxic control cells did not deviate significantly from uniformity (p = 0.673).

To quantify the extent of aerotactic migration across these different subpopulations, we computed the Forward Migration Index (FMI), a measure of directional persistence along a specified axis (Fig. 3A).^40,41^ The Parallel FMI (‖ FMI) captures displacement along the O_2_ gradient, where positive values indicate forward migration toward O_2_ and negative values reflect movement away from oxygen (Fig. 3A).^40,41^ The Perpendicular FMI (⊥ FMI) measures displacement orthogonal to the gradient, with values near zero representing random or unbiased lateral movement (Fig. 3A).^40,41^ We defined aerotactic behavior based on three criteria that have been reported previously: (i) ‖ FMI under hypoxia must be significantly higher than ‖ FMI under normoxic conditions; (ii) ‖ FMI must be significantly higher than ⊥ FMI within the same hypoxic condition, with ⊥ FMI values close to zero; and (iii) both ‖ and ⊥ FMI values under normoxia should be near zero.

Our data satisfy all criteria; PACCs exhibited the most positive ‖ FMI values, indicating enhanced directional migration toward O_2_, as well as ⊥ FMI values centered around zero (Fig. 3C, D). Non-PACC cells also showed modest but significant positive ‖ FMI values, consistent with a weaker aerotactic response, and ⊥ FMI values centered around zero (Fig. 3C, D). In contrast, cells under normoxia displayed FMI values near zero in both directions, which is to be expected under the conditions of uniform oxygen in the control (Fig. 3C,D).

To address the potential critique that the enhanced aerotactic migration observed in PACC cells may be simply due to their larger size, we performed correlational analyses between cell size and both ‖ FMI and ⊥ FMI across all conditions using Spearman’s ρ. (Fig. 3E,F, Supplementary Fig. S4). We found no significant correlation between cell area and directional migration based on Spearman’s ρ, suggesting that cell size alone does not account for the observed aerotactic behavior.

### PACCs express increased levels of HIF-1αand RhoA

HIF-1αis a key regulator of the hypoxic response, promoting cellular invasion through the upregulation of target genes such as *Snail* and *ZEB1*. ^42,43^ Thus, to investigate whether HIF-1αexpression is elevated in polyaneuploid cancer cells (PACCs), we performed fixed-cell immunofluorescence microscopy on PACC and non-PACC populations after 16 hours of hypoxia, as well as on normoxic controls. Since cells were stained with DAPI in these experiments, PACCs were identified based on a nuclear area ≥2.5x that of the normoxic parental population (Supplementary Fig. S2), consistent with established definitions. ^14,31,32^ Under hypoxic conditions, cells displayed markedly increased HIF-1αintensity with prominent nuclear localization compared to the low levels observed under normoxia (Fig. 4A-C, E-H, J). Quantification of Integrated Density, a measure of total fluorescence intensity per cell, revealed significantly higher HIF-1αexpression in PACCs compared to non-PACCs and normoxic controls (Fig. 4K).^44^ To ensure that the increased expression was not artificially inflated due to the large size of PACCs, Mean Fluorescence Intensity (MFI), reflecting per-unit-area expression, was also quantified and found to be highest in PACCs (Fig. 4L), indicating that the observed increase is not merely due to cell size.^44^

**Figure 4.**
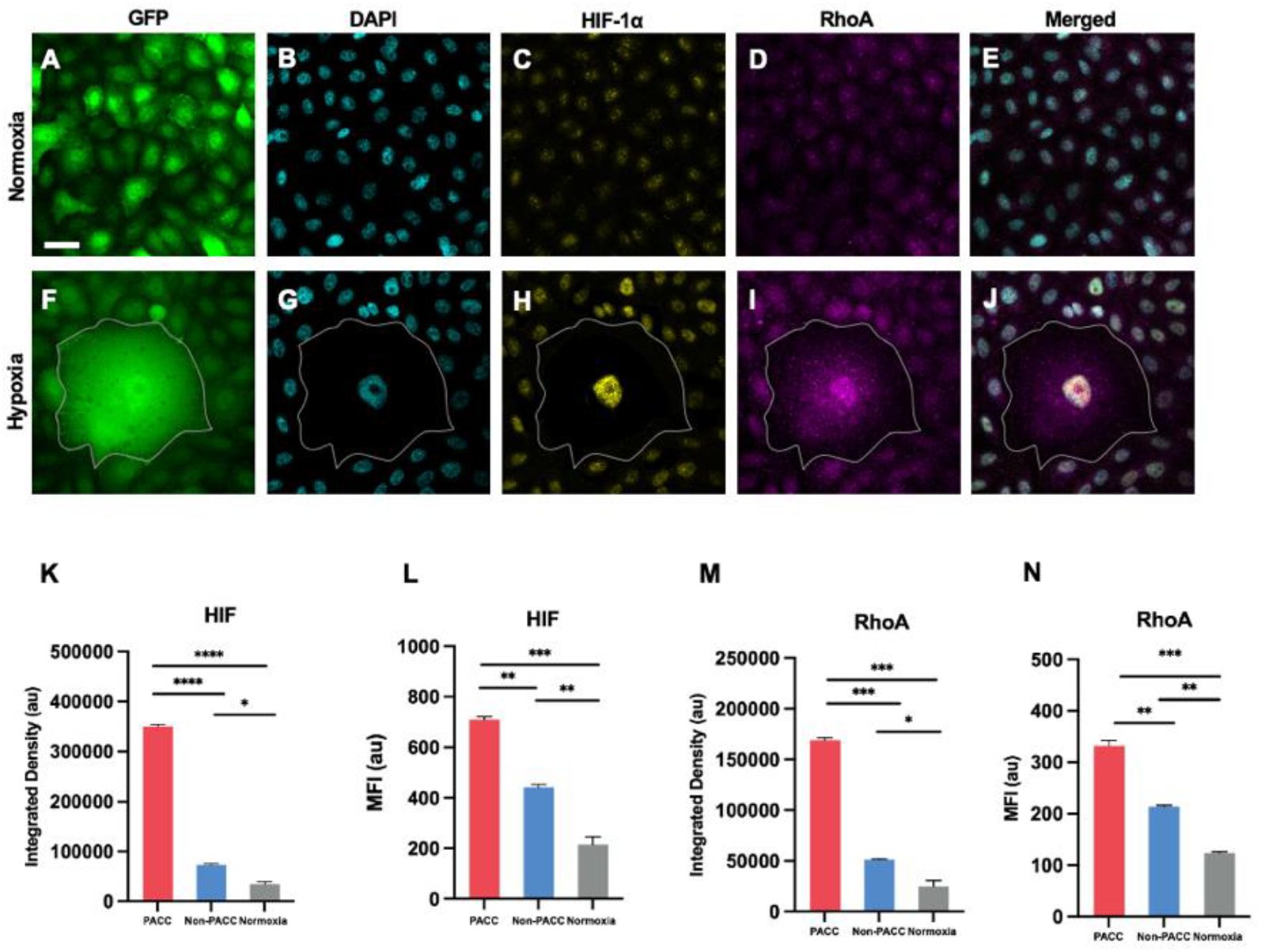
PACCs express increased levels of HIF-1α and RhoA. **(A-J)** Immunofluorescence microscopy of PC3 cells under normoxia **(A-E)** and hypoxia for 16 hours **(F-J)**, visualizing GFP **(A, F)**, DAPI **(B, G)**, HIF-1α (**C, H)**, and RhoA **(D, I).** Merged images **(E, J)** show GFP, DAPI, HIF-1α, and RhoA. PACCs under hypoxia (outlined in F-J) were identified by nuclear area ≥2.5x that of the normoxic parental population, as computed in Figure S2. **(K**,**L)** Quantification of HIF-1α intensity by Integrated Density **(K)** and Mean Fluorescence Intensity (MFI) **(L)** of PACC, Non-PACC, and Normoxic cells. **(M, N)** RhoA expression quantified by Integrated Density **(M)** and MFI **(N)** of PACC, Non-PACC, and Normoxic cells. Data represent mean ± SEM. Significance determined as p < 0.01 (^**^), and p < 0.001 (^***^) via One-way ANOVA with Tukey-Kramer post hoc test (see Materials and Methods). Scale bars for **A-J** are in **A** and represent 20 μm.

Given the role of RhoA in cytoskeletal remodeling and actomyosin contractility under hypoxia, we next assessed RhoA expression.^27–29,32^ RhoA was broadly cytoplasmic, with modest nuclear signal under hypoxia (Fig. 4D, E, I, J).

Similar to HIF1α, hypoxic cells exhibited significantly higher Integrated Density and MFI of RhoA staining compared to normoxic cells, with PACCs displaying the greatest RhoA expression among all groups (Fig. 4M, N).

### HIF-1αis required for the enhanced motility and aerotactic migration of PACC and non-PACC prostate cancer cells under hypoxia

To determine whether elevated HIF-1αexpression is necessary for the enhanced motility observed in PACCs, we depleted HIF1αusing siRNA-mediated knockdown and performed time-lapse fluorescence microscopy over 16 hours of hypoxia, as described above. HIF-1αknockdown was validated by quantifying MFI and Integrated Density in mock-transfected, control siRNA-transfected, and HIF-1αsiRNA-transfected cells that were fixed and stained after 16 hours of imaging (Fig. 5A-D). Depletion of HIF-1αresulted in cell trajectories that were notably shorter and less directed compared to cells treated with control siRNA (Fig. 5E-I). Quantification of motility metrics revealed significantly lower Accumulated and Euclidean distances, and Directness values closer to zero for both PACCs and non-PACCs following HIF-1αknockdown (Fig. 5J-L). These results suggest that the enhanced motility of hypoxic cells is HIF-1α-dependent.

**Figure 5.**
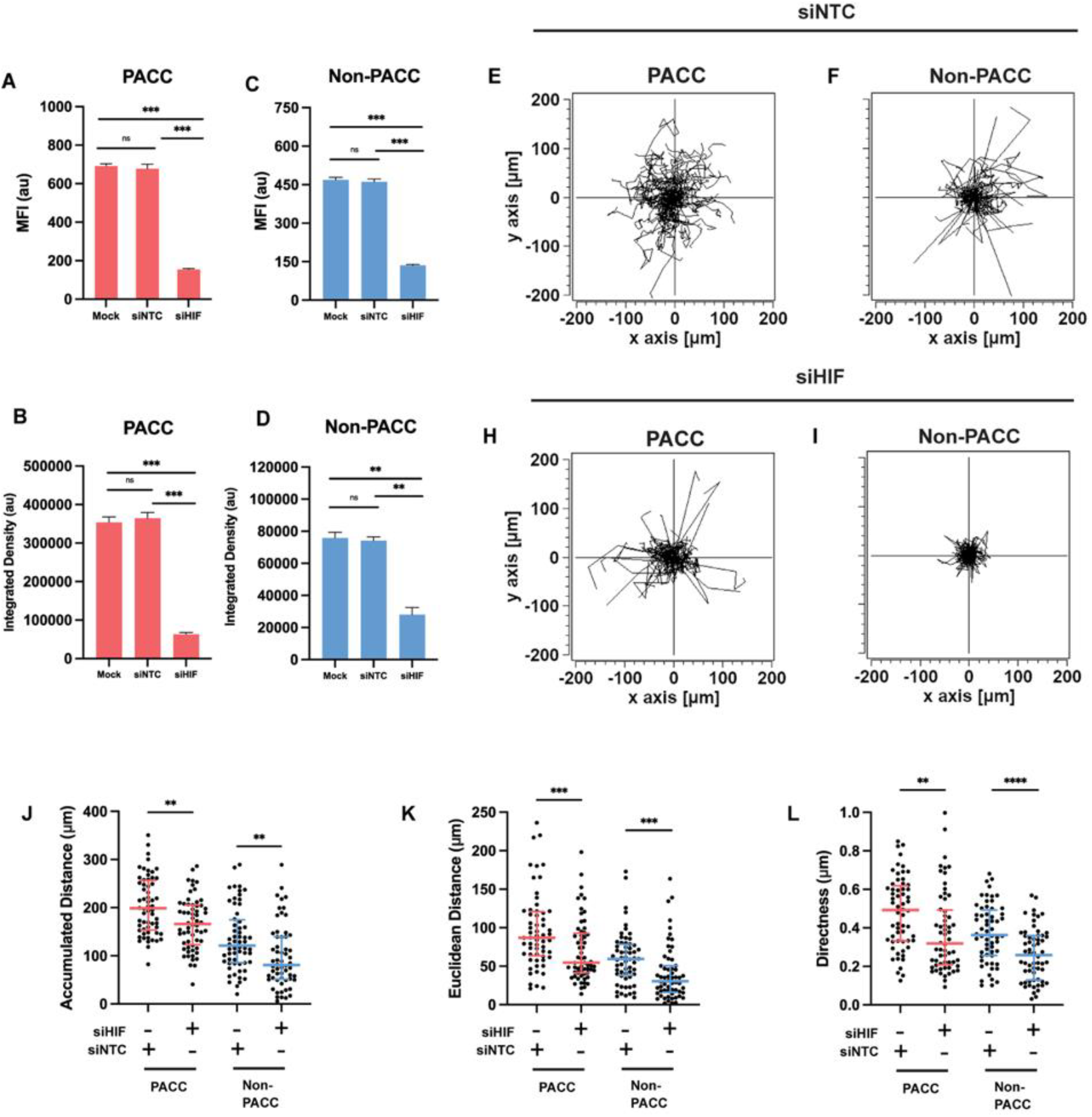
HIF-1α is required for the enhanced motility of PACCs, Non-PACC prostate cancer cells under hypoxia. (A–D) Validation of HIF-1α knockdown under hypoxia. Cells were mock transfected (Mock) or treated with non-targeting control siRNA (siNTC) or HIF-1α siRNA (siHIF). MFI and integrated density of HIF-1α staining were quantified in PACCs **(A, B)** and non-PACCs **(C, D)** after 16 hours of hypoxia. Data represent mean ± SEM. **(E–J)** Spider plots showing representative migration trajectories of PACCs and non-PACCs following control knockdown **(siNTC; E–G)** or HIF-1α knockdown **(siHIF; H–J**). n = 60 cells were sampled per group. **(K–M)** Quantification of Accumulated distance **(K)**, Euclidean distance **(L)**, and Directness **(M)** of tracked cells under hypoxia following siNTC or siHIF transfection. n = 60 cells per group. Data are shown as Median ± IQR. Significance determined by one-way ANOVA with Tukey-Kramer post hoc test in **(A–D)**, and by Mann-Whitney U test in **(K–M).** p < 0.01 (^**^), p < 0.001 (^***^), p < 0.0001 (^****^), ns = not significant.

We next sought to assess whether HIF-1αis required for the aerotactic migration observed in the hypoxic condition. The Rayleigh test showed no significant p values for the PACC and Non-PACC cells post-HIF1αknockdown (Fig. 6A). FMI analysis revealed that II FMI were significantly decreased for all three groups, with ⊥FMI remaining approximately zero in each group (Fig 6B-C). Altogether, these data suggest that HIF-1αis required for the aerotactic migration of prostate cancer cells under hypoxia.

**Figure 6.**
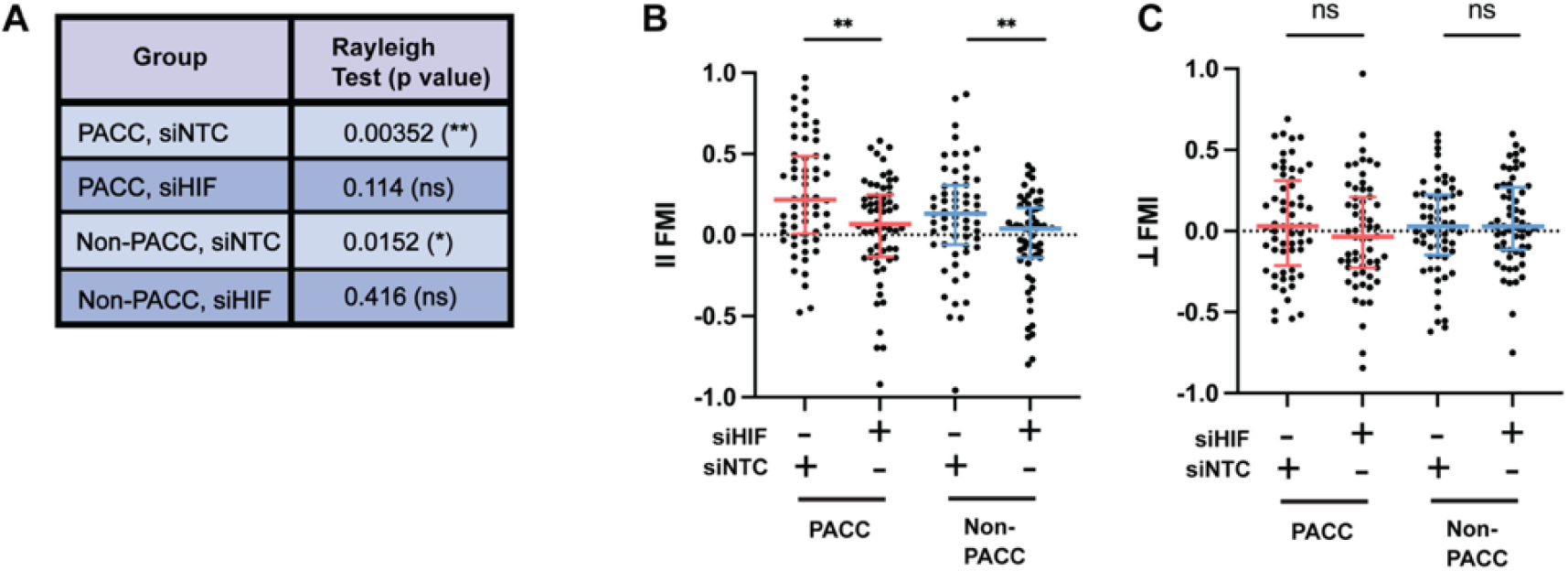
HIF-1α is required for the aerotactic migration of PACCs, Non-PACC prostate cancer cells under hypoxia. (**A)** Rayleigh test p-values for PACC and Non-PACC Cells following siNTC (control) or siHIF treatment (HIF-1α knockdown). **(B, C**) ‖ FMI **(B)** and ⊥ FMI **(C)** relative to the O_2_ gradient across cell types following siNTC or siHIF treatment. n = 60 cells per group. Data are shown as Median ± IQR. Significance in (**A)** was determined by the Rayleigh test. Statistical comparisons in **(B, C)** were made using the Mann-Whitney U test. p < 0.05 (*), p < 0.01 (^**^), and ns = not significant.

Importantly, under normoxic conditions, HIF-1αknockdown did not significantly alter the Accumulated Distance, Euclidean Distance, and Directness of prostate cancer cells, nor did it affect the II FMI or ⊥ FMI values. This suggests that HIF-1αis specifically required for hypoxia-driven migration, and not for migration under normoxia (Fig. 7).

**Figure 7.**
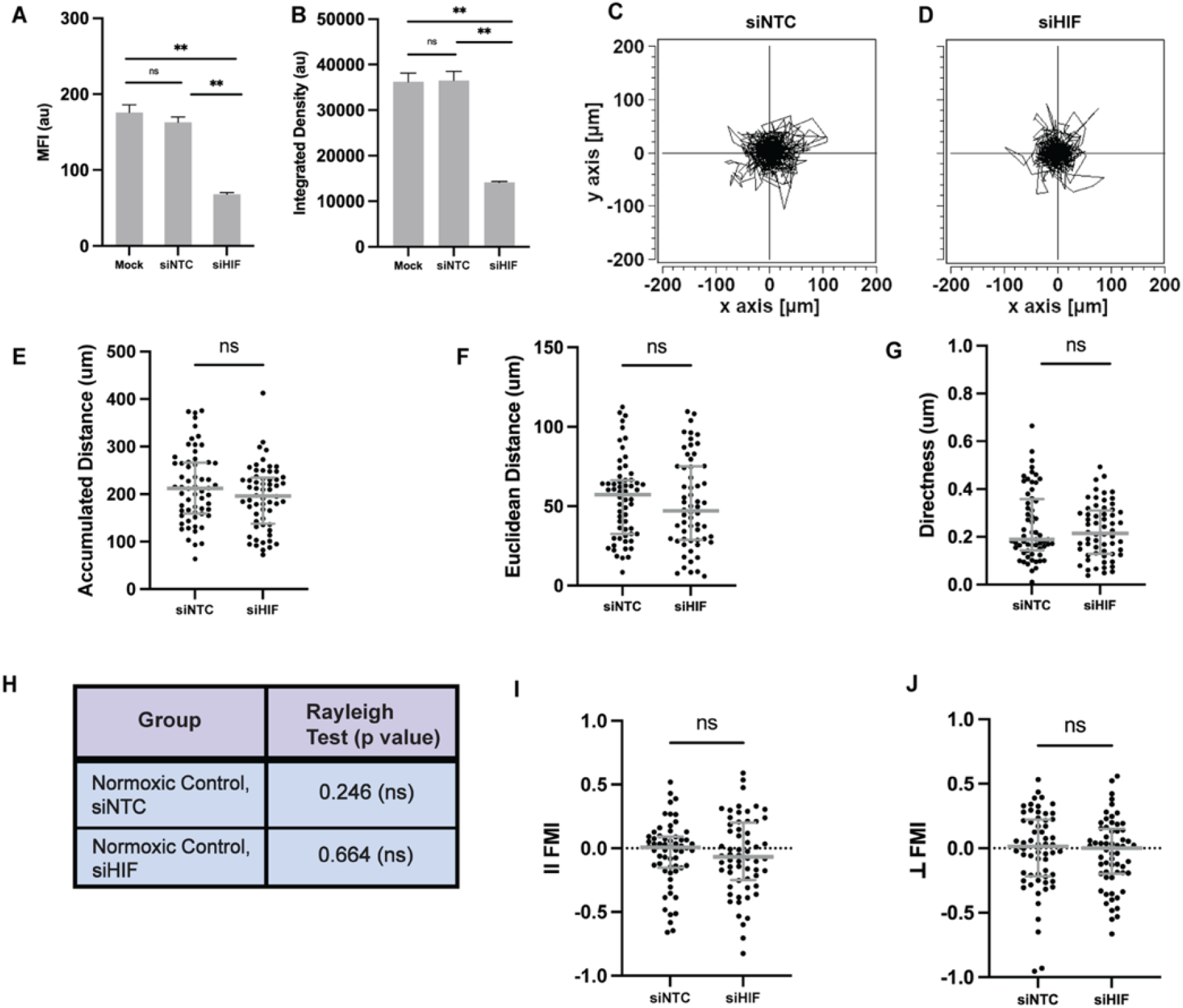
HIF-1α is not required for motility of prostate cancer cells in normoxia. **(A**,**B)** Validation of HIF-1α knockdown under normoxic conditions. Cells were either mock transfected (Mock), treated with non-targeting control siRNA (siNTC), or HIF-1α-targeting siRNA (siHIF). HIF-1α expression was quantified by measuring MFI **(A)** and Integrated Density **(B).** Data represent mean ± SEM. **(C, D)** Representative spider plots showing migration trajectories of normoxic cells following siNTC **(C)** or siHIF **(D)** treatment. n = 60 cells per group. **(E-G)** Quantification of Accumulated Distance (**E)**, Euclidean distance **(F)**, and Directness **(G)** of normoxic cells following siNTC or siHIF treatment (n = 60 cells per group). Data are shown as Median ± IQR. **(H)** Rayleigh test p-values in normoxic cells following transfection with siNTC or siHIF. **(I, J)** ‖ FMI **(I)** and ⊥ FMI **(J)** relative to the “O_2_ gradient” for normoxic cells following siNTC or siHIF treatment (n = 60 cells per group). Data are shown as Median ± IQR. Statistical significance was determined by one-way ANOVA with Tukey-Kramer post hoc test in **(A, B)**, Mann-Whitney U test in **(E–G, I, J)**, and Rayleigh test in **(H)**. p < 0.01 (^**^), ns = not significant.

### RhoA is required for the enhanced motility and aerotactic migration of PACC and non-PACC prostate cancer cells under hypoxia

Similar to HIF1α, we next depleted RhoA in PC3-GFP cells to assess whether the enhanced motility of PACCs is additionally RhoA-dependent. Time-lapse microscopy and validation of RhoA knockdown and were conducted as described for the HIF1αknockdown condition (Fig. 8A-D). RhoA depletion resulted in shorter and less directed cell trajectories compared to cells treated with control siRNA (Fig. 8E-I). This was supported by quantification of motility metrics: both Accumulated Distance and Euclidean Distance were significantly reduced, while Directness values decreased toward zero following RhoA knockdown (Fig. 8J-L). These results suggest that the increased motility of hypoxic cancer cells is also dependent on RhoA.

**Figure 8.**
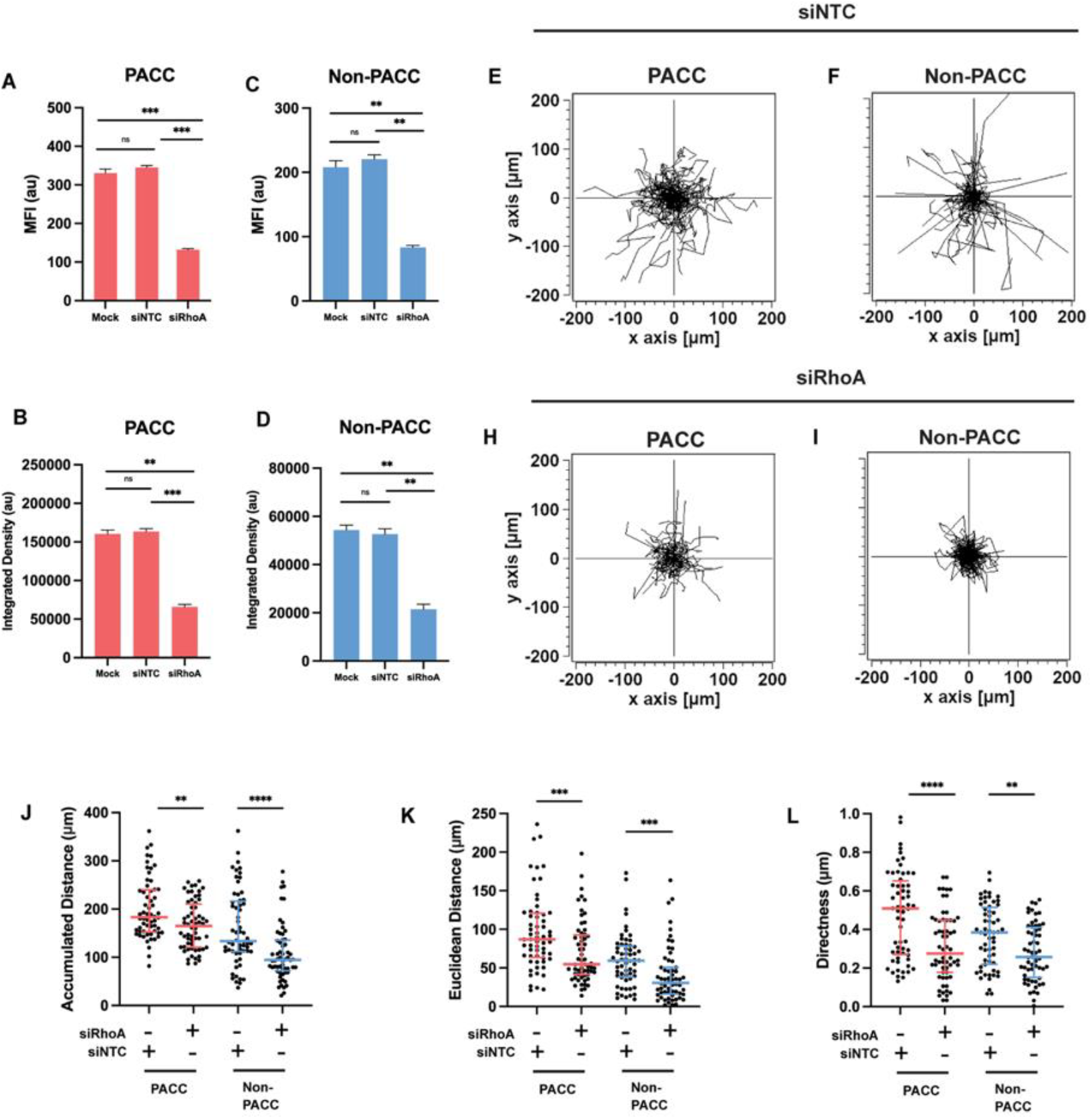
RhoA is required for the enhanced motility of PACCs, Non-PACC prostate cancer cells under hypoxia. **(A–D)** Validation of RhoA knockdown under hypoxia. Cells were mock transfected (Mock) or treated with non-targeting control siRNA (siNTC) or RhoA siRNA (siRhoA). MFI and integrated density of RhoA staining were quantified in PACCs **(A, B)** and non-PACCs **(C, D)** after 16 hours of hypoxia. Data represent mean ± SEM. **(E–J)** Spider plots showing representative migration trajectories of PACCs and non-PACCs following control knockdown (siNTC; **E-G**) or RhoA knockdown (siRhoA; **H-J**). n = 60 cells per group. **(K–M)** Quantification of Accumulated Distance **(K)**, Euclidean Distance **(L)**, and Directness **(M)** of tracked cells under hypoxia following siNTC or siRhoA transfection. n = 60 cells per group. Data are shown as Median ± IQR. Significance determined by one-way ANOVA with Tukey-Kramer post hoc test in **(A-D)**, and by Mann-Whitney U test in **(K-M).** p < 0.01 (^**^), p < 0.001 (^***^), p < 0.0001 (^****^), and ns = not significant

To assess whether RhoA is necessary for hypoxia-driven aerotaxis, we further quantified motility with respect to the O_2_ gradient. Similar to HIF-1αdepletion, Rayleigh tests post-knockdown yielded no significant p values for PACC and Non-PACC Cells (Fig. 9A). All three groups exhibited a significant decrease in II FMI relative that of their respective control knockdown condition (with ⊥ FMI remaining near zero) (Fig. 9B, C). These results suggest that RhoA is required for the aerotaxis of hypoxic prostate cancer cells.

**Figure 9.**
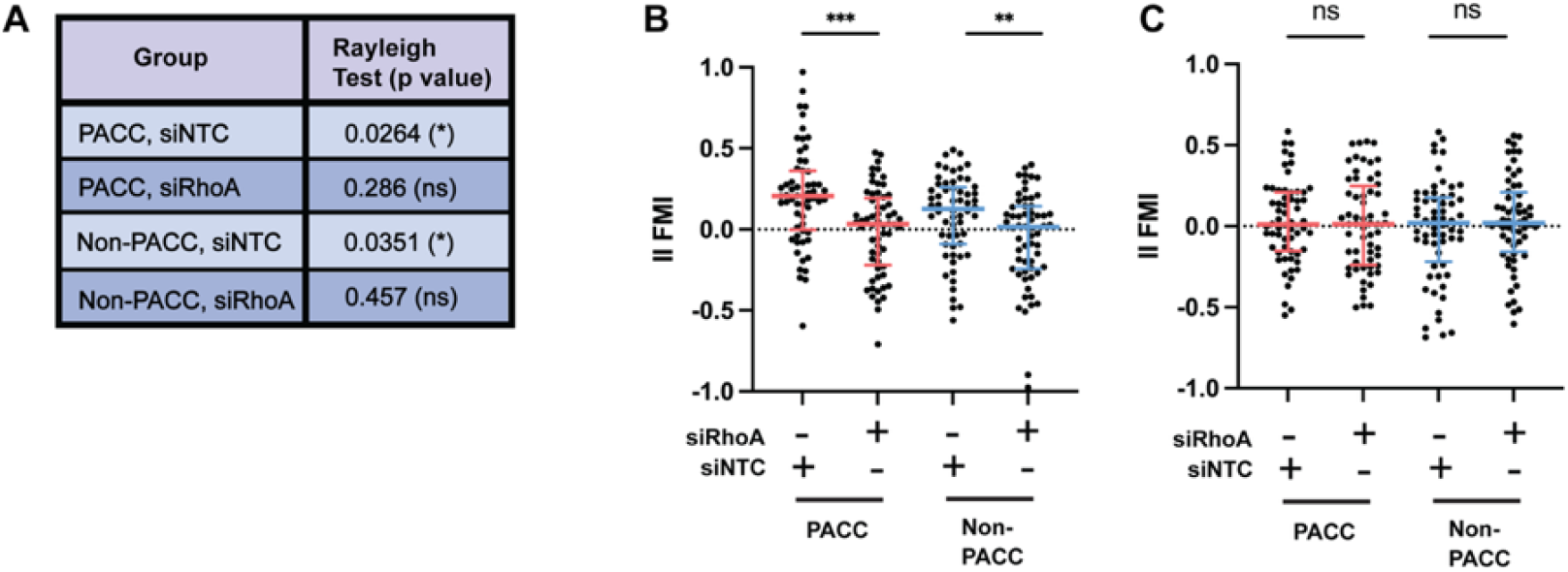
RhoA is required for the aerotactic migration of PACCs, Non-PACC prostate cancer cells under hypoxia. **(A)** Rayleigh test p-values for each group under hypoxia: PACC and Non-PACC Cells following siNTC or siRhoA treatment. **(B, C)** ‖ FMI **(C)** and ⊥ FMI **(D)** relative to the O_2_ gradient across cell types following siNTC or siRhoA treatment. n = 60 cells per group). Data are shown as Median ± IQR. Significance in **(A)** was determined by the Rayleigh test. Statistical comparisons in **(B, C)** were made using the Mann-Whitney U test. p < 0.05 (*), p < 0.01 (^**^), p < 0.001 (^***^), and ns = not significant.

Interestingly, under normoxic conditions, RhoA knockdown modestly reduced the Accumulated distance, Euclidean distance, and Directness of cells relative to control knockdown. No significant changes were observed in the II FMI or ⊥ FMI values, consistent with the lack of oxygen gradient present under normoxia (Fig. 10).

**Figure 10.**
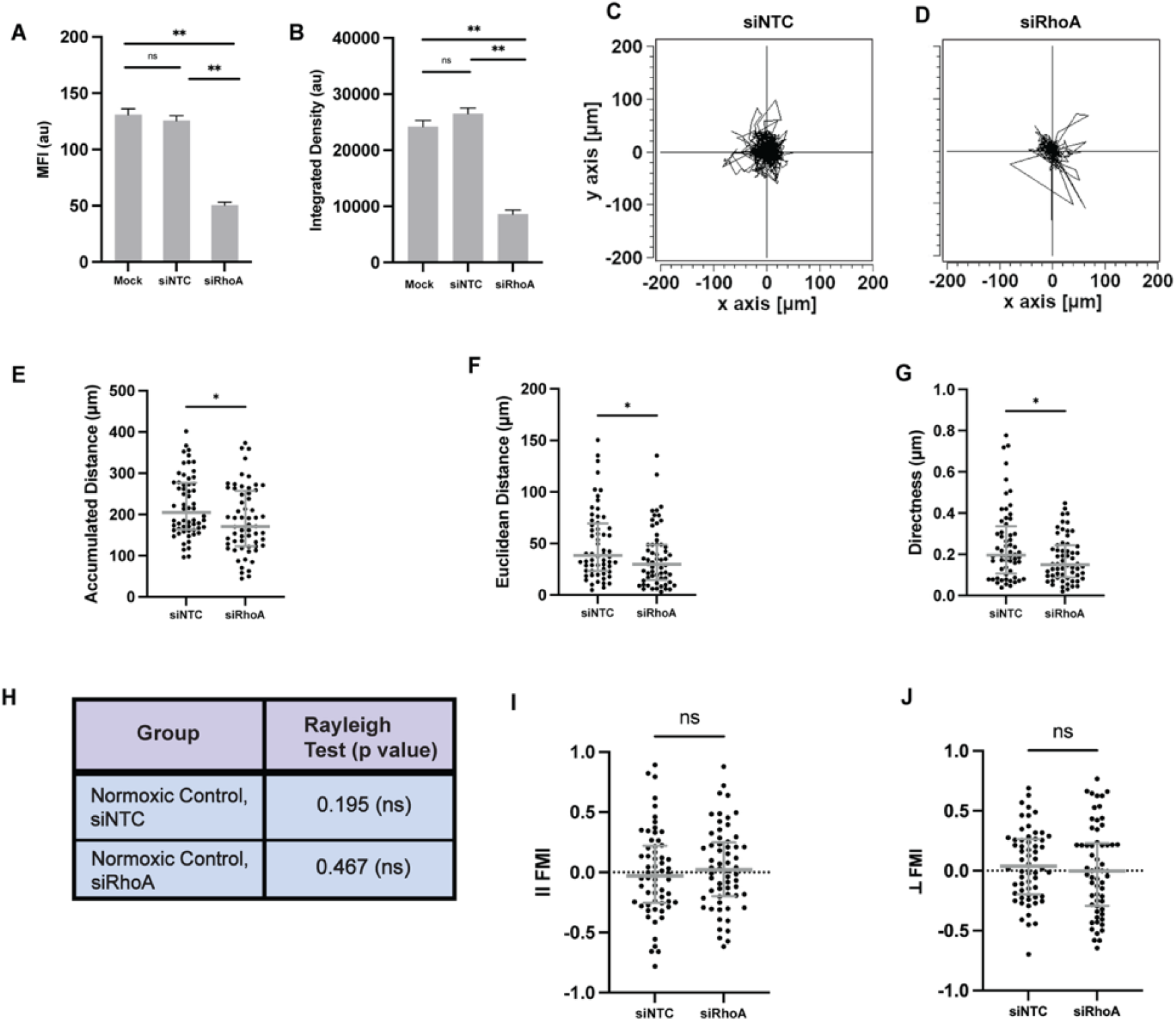
RhoA is required for the motility of prostate cancer cells in normoxia. **(A**,**B)** Validation of RhoA knockdown under normoxic conditions. Cells were either mock transfected (Mock), treated with non-targeting control siRNA (siNTC), or RhoA-targeting siRNA (siRhoA). RhoA expression was quantified by measuring MFI **(A)** and Integrated density **(B).** Data represent mean ± SEM. **(C, D)** Representative spider plots showing migration trajectories of normoxic cells following siNTC **(C)** or siRhoA **(D)** treatment. n = 60 per group. **(E–G)** Quantification of Accumulated Distance **(E**), Euclidean Distance **(F**), and Directness **(G**) in normoxic cells following siNTC or siRhoA treatment (n = 60 cells per group). Data are shown as Median ± IQR. **(H)** Rayleigh test p-values for biased migration in normoxic cells following transfection with siNTC or siRhoA. **(I, J)** ‖ FMI **(I)** and ⊥ FMI **(J)** relative to the “O_2_ gradient” for normoxic cells following siNTC or siRhoA treatment (n = 60 cells per group). Data are shown as Median ± IQR. Statistical significance was determined by one-way ANOVA with Tukey-Kramer post hoc test in **(A, B)**, Mann-Whitney U test in **(E–G, I, J)**, and Rayleigh test in **(H)**. p < 0.05 (*), ns = not significant.

Altogether, these results indicate that RhoA is necessary for hypoxia-induced motility and aerotactic migration, as well as for the motility of normoxic cells.

### RhoA expression is HIF-1αdependent in hypoxic prostate cancer cells

Previous studies in human colon cancer have reported that RhoA expression is HIF1α-dependent, while others have found that HIF-1αexpression is dependent on RhoA in renal cell carcinoma, highlighting a potentially cell type-specific directionality of the relationship between HIF-1αand RhoA.^29,30^ Since both HIF-1αand RhoA were found to be essential for the motility of PACC and non-PACC cells under hypoxic conditions, we next investigated whether a mechanistic link between them contributes to the enhanced migratory capacity observed in hypoxic prostate cancer cells, particularly in PACCs.

To determine whether HIF-1αis required for RhoA expression, we performed immunofluorescence microscopy as described previously to analyze RhoA fluorescence intensity levels in HIF-1α-depleted cells after 16 hours of hypoxia. Knockdown of HIF-1αresulted in a significant decrease in RhoA expression under hypoxia in both PACC and Non-PACC cells, and was supported by both MFI and Integrated Density quantification (Fig. 11A-Q). In contrast, normoxic PC3-GFP cells exhibited no significant changes in RhoA expression following HIF-1αknockdown (Fig. 12A-Q). These results suggest that RhoA expression in hypoxia is HIF-1α-dependent in prostate cancer cells, and that this regulatory relationship is specific to hypoxic conditions.

**Figure 11.**
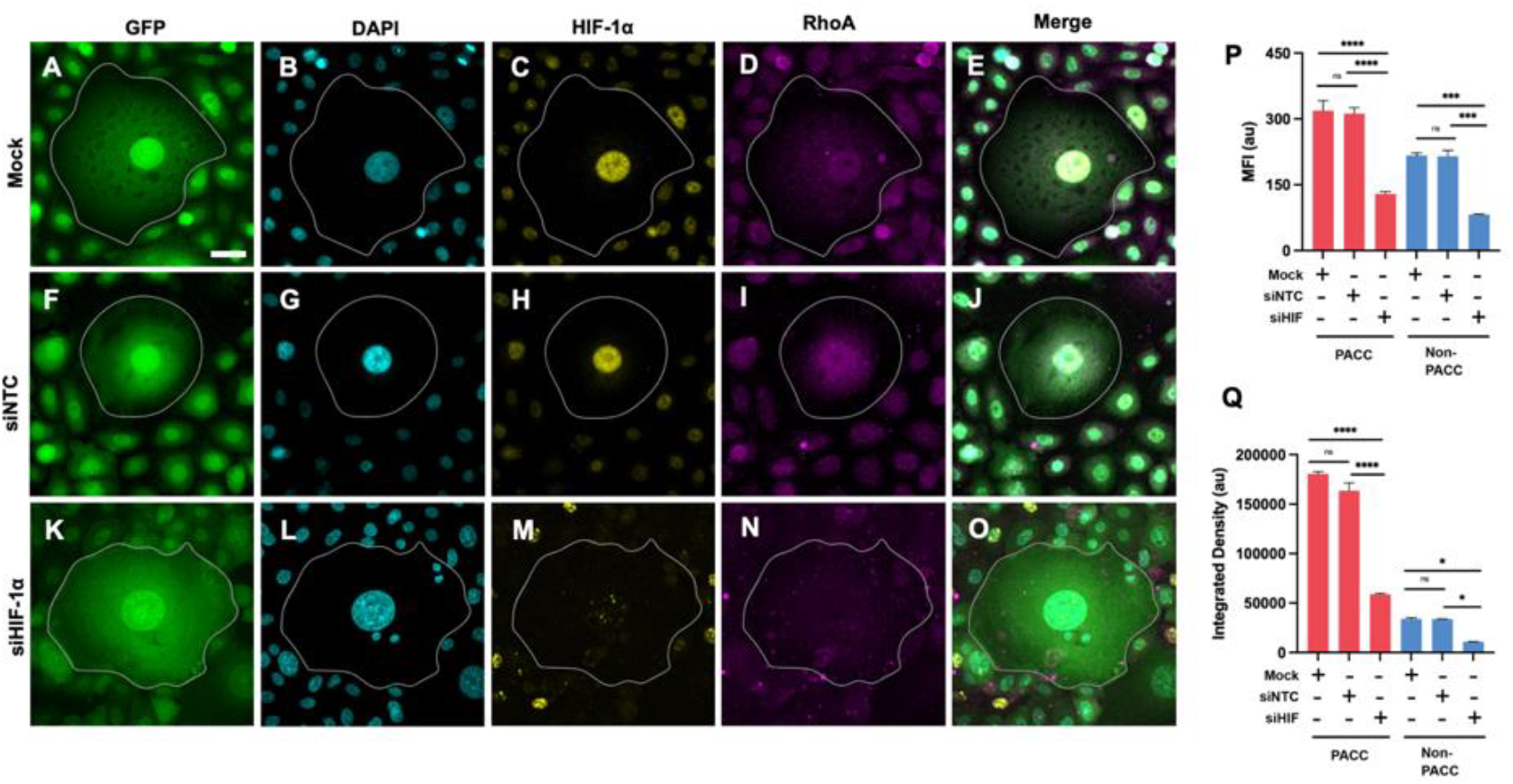
RhoA expression is HIF-1α dependent in hypoxic prostate cancer cells. **(A-O)** Immunofluorescence microscopy of PC3-GFP cells either mock transfected **(A-E)**, transfected with non-targeting control siRNA (siNTC, **F-J**), or transfected with HIF-1α siRNA (siHIF-1α, **K-O)** and exposed to hypoxia for 16 hours. Panels show GFP **(A, F, K)**, DAPI **(B, G, L)**, HIF-1α **(C, H, M)**, and RhoA **(D, I, N**). Merged images of all channels per condition are displayed in **(E, J, O).** PACCs (outlined in **A-O**) were identified by nuclear area ≥2.5x that of the normoxic parental population (see Fig. S2). **(P–Q)** Quantification of RhoA expression by MFI **(P)** and Integrated Density **(Q)** in PACC and non-PACC cells transfected with non-targeting control siRNA (siNTC), HIF-1α siRNA (siHIF), or mock transfected (Mock). Data represent mean ± SEM. Significance determined as p < 0.05 (*), p < 0.001 (^***^), p <0.0001 (^****^), and ns = not significant via One-way ANOVA with Tukey-Kramer post hoc test (see Materials and Methods). Scale bar in A represents 20 μm and applies to **A-O**.

**Figure 12.**
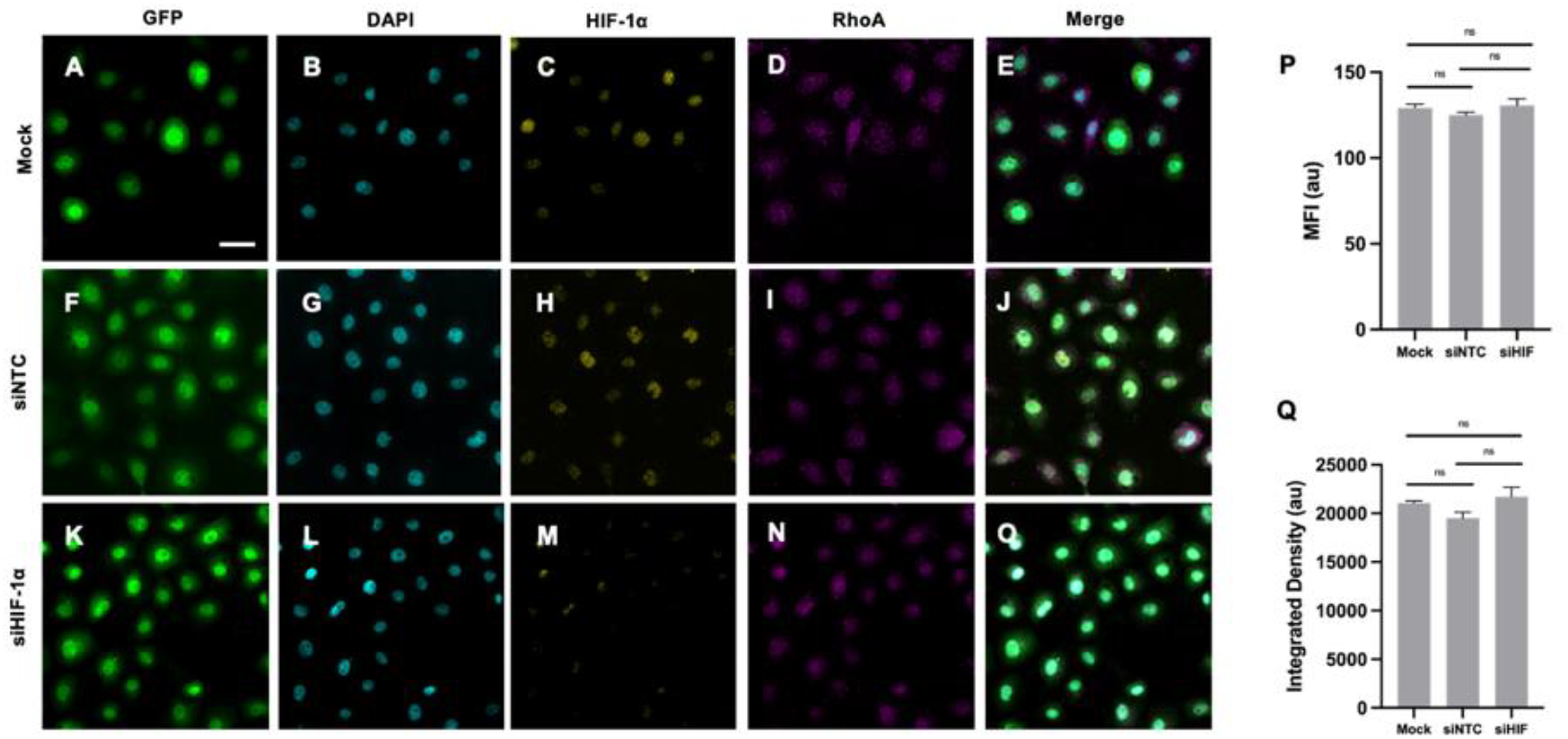
RhoA expression is not HIF-1α dependent in prostate cancer cells under normoxic conditions. **(A-O)** Immunofluorescence microscopy of PC3-GFP cells either mock transfected **(A-E)**, transfected with non-targeting control siRNA (siNTC, **F-J)**, or transfected with HIF-1α siRNA (siHIF-1α, **K-O)** under normoxic conditions. Panels show GFP **(A, F, K)**, DAPI **(B, G, L)**, HIF-1α **(C, H, M)**, and RhoA **(D, I, N**). Merged images of all channels per condition are displayed in **(E, J, O). (P–Q)** Quantification of RhoA expression by MFI **(P)** and Integrated Density **(Q)** in normoxic cells transfected with non-targeting control siRNA (siNTC), HIF-1α siRNA (siHIF), or mock transfected (Mock). Data represent mean ± SEM. Significance determined as ns = not significant via One-way ANOVA with Tukey-Kramer post hoc test (see Materials and Methods). Scale bar in A represents 20 μm and applies to **A-O**.

To further clarify the directionality of the relationship between HIF1αand RhoA, we assessed HIF1αexpression levels in RhoA-depleted PC3-GFP cells following 16 hours of hypoxia, as was done for HIF1α-depleted cells. Knockdown of RhoA resulted in no significant changes in the MFI or Integrated Density of HIF1α, either in PACC and Non-PACC cells under hypoxia, or under normoxic conditions (Supplementary Fig. S5A-Q, Supplementary Fig. S6A-Q). Taken together with the decreased RhoA expression observed following HIF1αdepletion, these results further suggest that, in prostate cancer, RhoA expression is dependent on HIF-1αunder hypoxic conditions, while HIF-1αexpression is independent of RhoA.

## DISCUSSION

Taken together, our results revealed that PACCs exhibited significantly increased motility and directional migration toward oxygen-rich regions under hypoxia. These behaviors were dependent on HIF-1α and RhoA, with evidence supporting a hypoxia-specific regulatory axis in which HIF-1α promotes RhoA expression.

Traditional *in-vitro* models used to study PACCs have often relied on extreme pharmacologic stressors (most notably high-dose cisplatin followed by FACS-based cell sorting) to purge proliferative cells and enrich for PACC populations.^9,10,31–33,45^ While these approaches have been instrumental in studies demonstrating the PACC state’s resistance and regenerative potential, they do not capture the cellular heterogeneity of tumors or accurately model the hypoxic microenvironments in which PACCs often arise. In contrast, our membrane-based culture system enables the self-generation of hypoxia by limiting O_2_ diffusion into the system’s core, thereby mimicking the O_2_ gradients that naturally emerge in poorly perfused tumor regions. FACS analysis revealed a nearly threefold enrichment of cells with >4N DNA content after 16 hours of hypoxia, consistent with previous reports showing a doubling of PACC frequency in patient-derived xenografts treated with the PARP inhibitor olaparib.^45^ These results suggest that our system not only recapitulates key features of the hypoxic tumor microenvironment, but also models important aspects of tumor heterogeneity. To further simulate the tumor niche, future studies may co-culture PC3 prostate cancer cells with human stromal fibroblasts under hypoxic conditions. Notably, recent research shows that hypoxia drives a pro-inflammatory phenotype in cancer-associated fibroblasts (CAFs), leading to the secretion of chemokines that promote cancer cell migration, invasion, and the emergence of cancer stem cells (CSCs).^46–48^ Given the functional similarities between CSCs and PACCs, particularly their shared regenerative potential, it is plausible that CAFs may also promote entry into the PACC state – a possibility that remains unexplored and warrants future investigation.

Single-cell tracking within our culture system revealed that PACCs exhibit significantly increased total and directional motility compared to both non-PACCs and similarly sized large cells that do not undergo rapid expansion. Additionally, PACCs demonstrated the most robust forward migration along O_2_ gradients generated by this system, suggesting enhanced aerotactic migration. These findings support the idea that PACCs may be particularly well equipped to complete key steps of the metastatic cascade. Indeed, tumor cells must not only survive hypoxic stress but also migrate toward oxygenated blood vessels to escape the primary tumor and initiate dissemination.^8,34^ Thus, the observed motility and aerotactic behavior suggest that PACCs may be especially well suited to facilitate invasion and intravasation.

Interestingly, an often-overlooked interpretation of aerotactic migration is its potential role in tumor escape rather than just systemic spread.^22^ Prior studies in breast tumor cells have suggested that directional migration toward O_2_ -rich regions may enable escape from the most hypoxic zones within the tumor itself.^22^ In this context, aerotaxis may not only serve as a predictor of metastatic potential but may also represent a survival mechanism that supports PACC persistence within the tumor microenvironment.^22^ Future studies using our membrane-based culture system may aim to directly assess PACC survival under hypoxia by incorporating cell death markers into our analysis.

This study additionally investigates the molecular mechanisms by which PACCs may acquire their enhanced motility and aerotactic behavior. While HIF-1αis widely recognized as a master regulator of genes involved in cellular adaptation to hypoxia – including *VEGF*, which promotes angiogenesis, and *GLUT1*, which facilitates anaerobic metabolism – its role in regulating aerotactic migration has not been well characterized.^24,49,50^ We observed upregulation of HIF-1α under hypoxic conditions, particularly in PACCs, and found that knockdown of HIF-1α led to a marked reduction in both overall motility and directional migration toward oxygen. These findings suggest a previously underappreciated role for HIF-1α in regulating aerotaxis, extending its influence beyond metabolic and angiogenic adaptation to include directional migration in response to microenvironmental O_2_ gradients.

Similarly to HIF-1α, we observed upregulation of the small GTPase RhoA under hypoxic conditions, with the highest expression levels observed in PACCs. Knockdown of RhoA significantly reduced both total and net distances traveled, as well as the directness of cell movement and aerotactic migration in hypoxic cells. Interestingly, RhoA knockdown also led to a modest but statistically significant decrease in motility under normoxic conditions. These findings suggest that while RhoA contributes to cancer cell motility in both environments, its role may be particularly critical under hypoxic stress.

Furthermore, siRNA-mediated knockdown of HIF-1α and RhoA supports an interplay between these two proteins under hypoxic conditions. To our knowledge, this is the first study to suggest a regulatory relationship between HIF-1α and RhoA in prostate cancer. Specifically, depletion of HIF-1α led to a significant decrease in RhoA expression under hypoxia, whereas no significant change was observed under normoxic conditions. In contrast, RhoA knockdown did not significantly alter HIF-1α levels in either condition, further suggesting that RhoA expression is HIF-1α -dependent, particularly in hypoxia. Altogether, we propose a mechanism by which directed motility and aerotactic migration are upregulated in PACCs, potentially via a HIF-1α -RhoA signaling axis. In our model, intratumoral hypoxia stabilizes HIF-1α, leading to the transcriptional activation of RhoA. RhoA, in turn, promotes actin polymerization and actomyosin contractility likely through the canonical RhoA–ROCK1/2 pathway, facilitating cytoskeletal reorganization, directed cell migration, and aerotaxis.

More broadly, our findings suggest that PACCs may represent a therapeutically relevant subpopulation, characterized by enhanced motility and oxygen-sensing capacity under hypoxia. We identify HIF-1α and RhoA as key regulators of these behaviors, both of which are amenable to small-molecule inhibition.^51,52^ Targeting these proteins may offer a strategy to disrupt the metastatic progression and therapeutic resistance associated with PACCs.

### Limitations of the study and more future directions

Our findings highlight new opportunities to further refine our understanding of PACC dynamics. Notably, our single-cell tracking experiments did not incorporate nuclear staining and instead correlated nuclear content from cell size (Fig. 2C, 2D, S2). This approach was taken as commonly used DNA-binding dyes are reported to be suboptimal for long-term live-cell imaging due to their potential interference with replication and transcription, and UV-induced phototoxicity.^35–37^ To enhance the precision of PACC identification in future live-cell imaging studies, future work may incorporate Fluorescent Ubiquitination-based Cell Cycle Indicators (FUCCI), which enables real-time, non-invasive monitoring of cell cycle progression.^53^

Furthermore, future work should aim to more precisely quantify the expression of PACCs by incorporating cell volume measurements in addition to the 2D cell area used in this study. Z-stacks enable accurate measurement of cell depth, which can then be used to calculate cell volume – an important consideration given the enlarged morphology of PACCs. In this context, Z-stacks also allow for more reliable quantification of protein expression by summing fluorescence intensities across multiple planes, rather than relying solely on the brightest focal plane.^54^ This approach will provide a more comprehensive understanding of PACC-specific protein expression and its relationship to cell morphology under hypoxic conditions.

## METHODS

### Cell culture

All experiments were performed with the PC3-GFP prostate cancer cell line kindly provided by the Pienta laboratory. The PC3-GFP cell line was originally generated by labeling standard PC3 cells (ATCC, CRL-1435) with the pLenti-CMV-GFP-Neo (657-2) GFP expression vector (Addgene, Plasmid 17447). Cells were cultured at 37 degrees Celsius in 5% CO_2_ in RPMI 1640 media (Gibco, Thermofisher) supplemented with 10% Fetal Bovine Serum (Thermo Fisher Scientific) and 1% Penicillin/Streptomycin (Gibco, Thermofisher). Authentication by STR profiling and routine mycoplasma testing were performed by the Pienta laboratory. All cells used in this study were confirmed to be mycoplasma-free.

### siRNA transfection

PC3-GFP cells were transfected with siRNA using Lipofectamine RNAiMAX (Invitrogen) in Opti-MEM (Gibco) without antibiotics. Medium was changed to RPMI 1640 media (Gibco, Thermofisher) with 10% Fetal Bovine Serum (Thermo Fisher Scientific) and 1% Penicillin/Streptomycin (Gibco, Thermofisher) 6 hours after transfection. Cells were analyzed 48 hours post-transfection.

The following siRNAs from Dharmacon (Thermo Fisher Scientific) were utilized: ON-TARGETplus Human HIF1A (3091), ON-TARGETplus Human RhoA (387) siRNA-SMARTpool, ON-TARGETplus Non-targeting Pool siRNA-SMARTpool, and Accel eGFP Pool.

### Experimental setup for self-generated hypoxia

Hypoxia was induced using our previously described membrane-based system (Hosny et al.), which establishes a stable oxygen gradient across the dish. In brief, cells were seeded at a density of 0.8 × 10^6^ cells per 60 mm Lumox gas-permeable dish and cultured to approximately 80% confluence. A phosphorescent thin film composed of platinum porphyrin (PtTFPP) embedded in a perfluoropolyether (PFPE) matrix (kindly provided by the Qu Lab) was mounted on a glass coverslip with the film side placed against the bottom of the Lumox dish. Since the red PtTFPP phosphorescence is quenched in the presence of oxygen, the intensity signal enabling us to monitor and confirm hypoxia development in real time.

A 30 mm diameter acrylic plug was then carefully positioned over the center of the dish above the cells. Micropillars (100 µm in height) on the plug base minimize media flow that could displace cells, while the uniform spacing ensure cells remained uncompressed throughout the experiment Following plug placement, the dish was transferred to an on-stage CO_2_ incubator (OkoLab) at 37°C, and cells were tracked during hypoxia development over 16 hours.

### Time-lapse microscopy and single-cell tracking

Tiled images of PC3-GFP cells (GFP channel) and the phosphorescent thin film (mCherry channel) were taken every 0.5 hours over the course of hypoxia development using the Nikon Eclipse TE2000-E inverted microscope. Images were acquired with a Plan Fluor 4X objective and an Andor Zyla cooled camera, and all hardware was controlled using NIS Elements software.

We used the Cellpose module within the TrackMate plugin in Fiji (Version 1.52p) to segment the time-lapse images.^55,56^ Cell detection was performed using the *cyto2* pretrained model.^56^ The resulting data were further analyzed in TrackMate, and position and cell area measurements were acquired at each time point.

### Cell Motility Measurements

Time-lapse position data were exported for quantification in R software. Three parameters were computed to assess cell motility: Accumulated Distance, Euclidean Distance, and Directness.

Accumulated Distance (*D*_*accumulated*_) was calculated as the sum of the distances between consecutive cell positions (*x, y*) across all time points, where *x*_*i*_ and *y*_*i*_ represent the cell’s *x*- and *y*-coordinates at time point *i*, and *n* is the total number of time points:

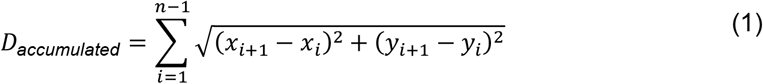

Euclidean Distance (*D*_*euclidean*_) represents the straight-line distance between the starting and ending cell positions (i.e., the net distance traveled), where (*x*_1_, *y*_1_) are the coordinates of the cell at the initial time point, (*x*_*n*_, *y*_*n*_) are the coordinates at the final time point:

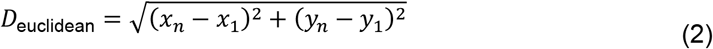

Directness was computed as the ratio of *D*_*euclidean*_ to *D*_*accumulated*_:

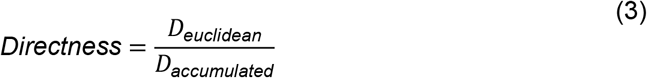

To normalize data and account for cells that were unable to be tracked for the full 16-hour hypoxia experiment, trajectories were scaled by multiplying all *x* and *y* positions by a factor 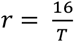, where *T* represents the actual duration of the trajectory in hours. Cells with fewer than 3 frames (<1 hour of tracking data) were excluded from analysis. A total of 60 cells were analyzed per condition.

### Forward Migration Index (FMI) Calculations

For subsequent calculations involving the parallel and perpendicular components of the forward migration index (*FMI*_∥_ and *FMI*_⊥_, respectively) of cell trajectories, the O_2_ gradient was approximated as a vector field pointing radially outward from the center of the system. Specifically, this gradient vector ***g*** = (*g*_*x*_ *g*_*y*_)^*T*^ = ∇φ was assumed to point from the system center to the midpoint of each trajectory, which was obtained by averaging the initial and final (*x, y*) coordinates of each individual cell.

Given a displacement vector

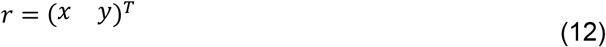

the parallel and perpendicular components of the trajectory with respect to the gradient vector are, respectively, given by:

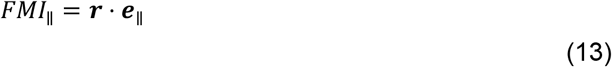

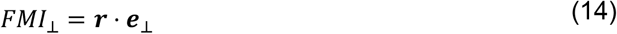

where the parallel and perpendicular unit vectors corresponding to the O_2_ gradient ***g*** = (*g*_*x*_ *g*_*y*_)^*T*^ = ∇φ are

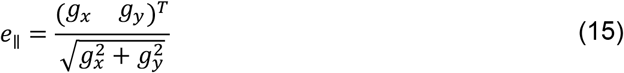

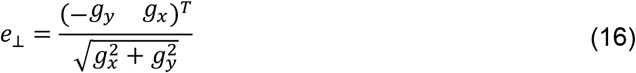

These computations were conducted in R software (version 4.5.0).

### Spider Plots

Cell trajectory data from TrackMate were imported into the Chemotaxis and Migration Tool for visualization in spider plots, which display the (x, y) coordiantes of the trajecorties with initial positions mapped to the origin (0,0).

### Immunofluorescence

Following 16 hours of hypoxia, the acrylic plug was removed and cells were immediately fixed in 4% paraformaldehyde (PFA) (Thermo Fisher Scientific) for 15 minutes. After fixation, cells were washed five times for five minutes each using 0.05% Triton X-100 Surfact-Amps in PBS (Thermo Fisher Scientific). Blocking was performed for 1 hour at room temperature using a solution of 5% Normal Goat Serum (Abcam) in PBS. Cells were then incubated overnight at 4 °C with the primary antibody, followed by five additional 5-minute washes in 0.05% Triton X-100 in PBS. Secondary antibody incubation was carried out for 1 hour at room temperature. Afterward, cells were washed again five times for five minutes each. Nuclear staining was performed using a 1X solution of PureBlue DAPI Nuclear Staining Dye (Bio-Rad, #1351303) for 15 minutes, followed by three final washes (5 minutes each). PBS was then added, and the samples were imaged using a confocal microscope.

### Antibodies for Immunostaining

Primary antibodies were diluted to a concentration of 1:100: rabbit anti-HIF1 alpha (Abcam, ab51608), mouse anti-RhoA (Proteintech, 66733-1-IG). Secondary antibodies were diluted to a concentration of 1:500: Goat anti-Rabbit IgG (H+L) Highly Cross-Adsorbed Alexa Fluor 555 (Invitrogen, A32732), Goat anti-Mouse IgG (H+L) Highly Cross-Adsorbed Alexa Fluor 647 (Invitrogen, A21236). All antibodies were diluted in a solution of 1% Bovine Serum Albumin (BSA) Fraction V (Thermo Fisher Scientific), 0.05% Triton X-100 Surfact-Amps, 0.5% Normal Goat Serum in PBS.

### Confocal Imaging

Tiled images of immunostained cells were acquired at 20X magnification on the Nikon Ti-E Inverted Microscope equipped with TIRF, Super-Resolution (localization type), and Ablation modules (TIRF/DISC2 system). The 405 nm (to visualize DAPI), 488 nm (GFP), 561 nm (HIF1α), and 640 nm (RhoA) laser lines were used. Image acquisition was performed using NIS-Elements software. All images were acquired using the same laser power, exposure time, and gain per channel across experimental repeats: 640 nm (900 ms, 18% power, gain 1.0); 561 nm (1s, 60% power, gain 1.0), 488 nm (400 ms, 30% power, gain 1.0); 405 nm (300 ms; 34% power, gain 1.0).

### Quantification of Immunofluorescence Microscopy

Multi-channel immunofluorescence image analysis was performed using the TrackMate/Cellpose plug-in described above.^55,56^ To quantify Mean Fluorescence Intensity (MFI) per cell, individual cells were segmented separately in each channel using the *cyto 2* pretrained model.^56^ The position of each cell, cell area, and raw MFI values were exported for further analysis in R. To match cells across channels, the Fast Nearest Neighbor (FNN) R package was used to identify corresponding cells based on their proximity in (x, y) coordinates within each channel. MFI values were manually background-subtracted using the intensity of a nearby cell-free region acquired in Fiji. Background-subtracted MFIs and TrackMate-derived cell areas were then used to compute Integrated Density for each cell (MFI x Cell Area).

### FACS-Based Identification of PACCs

Flow cytometry was performed on ∼70,000 hypoxic and ∼70,000 normoxic cells per replicate using the FACSymphony A3 flow cytometer (BD Biosciences). Immediately post-hypoxia, cells were harvested with TrypLE Express (Gibco), fixed in 4% PFA on ice for 15 minutes, centrifuged and washed twice with PBS. Cells were then stained with a 1X solution of DAPI for 15 minutes, followed by two additional PBS washes.

Doublets were excluded via forward scatter area versus height (FSC-A vs. FSC-H) and DAPI width versus area (UV450-W vs. UV450-A) gating strategies. DAPI was excited with the UV laser at 355nm and emitted fluorescence was collected through a 450/50nm bandpass filter. Data were analyzed using FlowJo software (FlowJo, LLC), and cells with >4N DNA content based on DAPI intensity were classified as PACCs.

### Statistics and Reproducibility

All statistical analysis was conducted with Prism 9. Data was tested for normality using the Shapiro-Wilk Test to determine appropriate statistical tests. For motility and aerotaxis measurements (Accumulated Distance, Euclidean Distance, Directness, and FMIs), pairwise comparisons of n = 60 cells per group were performed using non-parametric T-tests (Mann-Whitney U Tests), while multiple group comparisons were analyzed using Kruskal-Wallis tests followed by Dunn’s post hoc test. Quantification of flow cytometry (Fig. 3I) and nuclear area fold change vs cell area scatterplot data (Fig. 4D) was performed using Welch’s t-test. One-way ANOVA with Tukey-Kramer post hoc test was used to analyze all immunofluorescence MFI and Integrated Density data. Correlation analyses between cell area and all motility and aerotaxis parameters were conducted using Spearman’s rank correlation coefficient (Spearman’s ρ). Significance was determined via the following: ns = not significant, p < 0.05 (*), p < 0.01 (**), p < 0.001 (***), p < 0.0001 (****).

## Data availability

The datasets generated during and/or analyzed during the current study are available from the corresponding author on reasonable request.

## Acknowledgements

This work was supported by the Johns Hopkins University Discovery Award 2023– 2024, US National Science Foundation (PHY-1659940 and PHY-1734030), the National Natural Science Foundation of China (2421003 and 62127819), and the Class of 1955 Senior Thesis Fund for Fall 2024. Fig. 2E and 3A were created using BioRender.

