## Supplemental Figures S1-S6 for "HIF-1α and RhoA Drive Enhanced Motility and Aerotaxis of Polyaneuploid Prostate Cancer Cells in Hypoxia"

**Supplementary Figure S1. Cell cycle distributions of PC3-GFP cells under hypoxic and normoxic conditions**. **(A)** Representative DNA content histogram of PC3-GFP cells under normoxic conditions shows typical distribution across G1, S, and G2/M phases, with a small proportion (2.70%) of cells exhibiting DNA content greater than 4N **(B)** DNA content histogram of PC3-GFP cells in hypoxia reveals a marked increase in the >4N population (7.20%). Insets in **(A,B)** highlight the >4N population in normoxia and hypoxia, respectively.

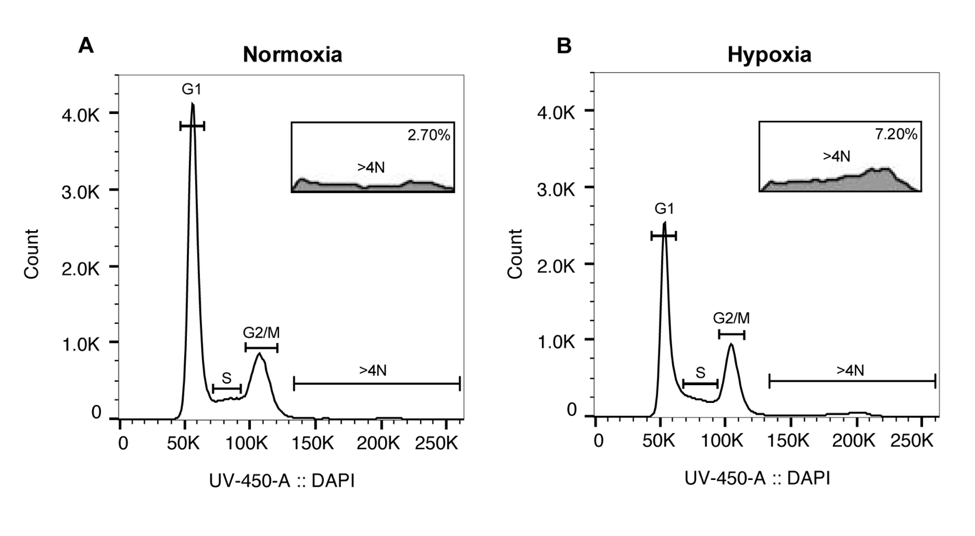

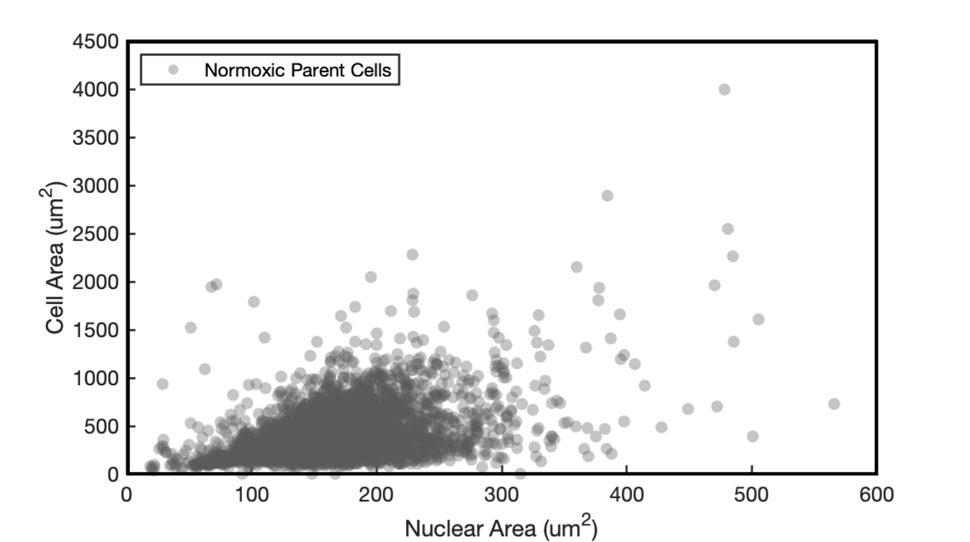

**Supplementary Figure S2: Nuclear area versus cell size scatterplot of normoxic parent cells. (A)** Normoxic parent cells exhibited a mean nuclear area of 165 µm^2^ ±  0.64 µm^2^ (mean ± SEM) and mean cell area of 403 µm^2^ ± 3.2 µm^2^ (mean ± SEM).

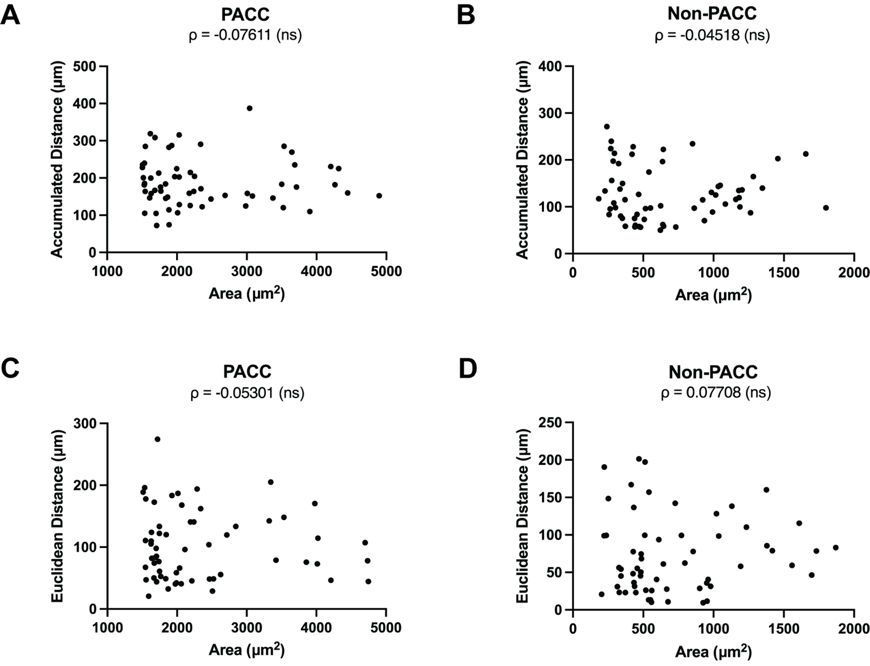

**Supplementary Figure S3: Cell size versus motility measurements. (A-F)** Scatterplots of Accumulated Distance vs Cell Area **(A-C)** and Euclidean Distance vs Cell Area (**D-F)** including Spearman's ρ for PACC and Non-PACC Cells.

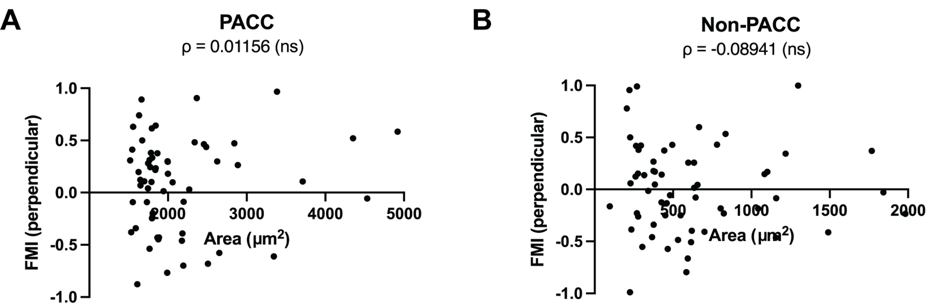

**Supplementary Figure S4: Cell size versus aerotaxis measurements. (A-C)** Spearman's ρ and Scatterplots of Perpendicular FMI vs Cell Area for PACC **(A)** and Non-PACC **(B)** Cells.

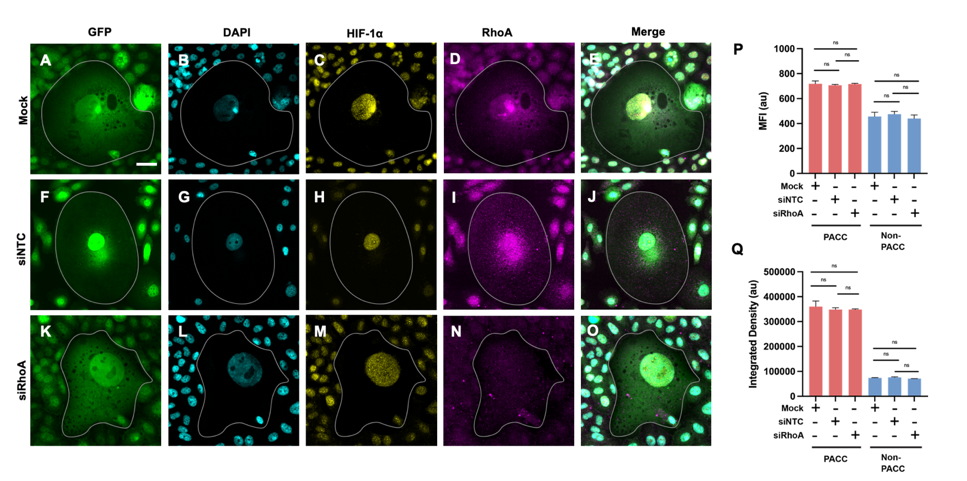

**Supplementary Fig. S5. HIF-1α expression is independent of RhoA in hypoxia. (A-O)** Immunofluorescence microscopy of PC3-GFP cells either mock transfected **(A-E),** transfected with non-targeting control siRNA **(**siNTC**, F-J**), or transfected with RhoA siRNA **(**siRhoA**, K-O)** and exposed to hypoxia for 16 hours. Panels show GFP **(A, F, K),** DAPI **(B, G, L)**, HIF-1α **(C, H, M)**, and RhoA **(D, I, N**). Merged images of all channels per condition are displayed in **(E, J, O).** PACCs **(**outlined **in A-O)** were identified by nuclear area ≥2.5x that of the normoxic parental population (see Fig. S2). **(P–Q)** Quantification of HIF-1α expression by MFI **(P)** and Integrated Density **(Q)** in PACC and non-PACC cells transfected with non-targeting control siRNA (siNTC), RhoA siRNA (siRhoA), or mock transfected (Mock). Data represent mean ± SEM. Significance determined as ns = not significant via One-way ANOVA with Tukey-Kramer post hoc test (see Materials and Methods). Scale bar in **A** represents 20 μm and applies to **A-O**.

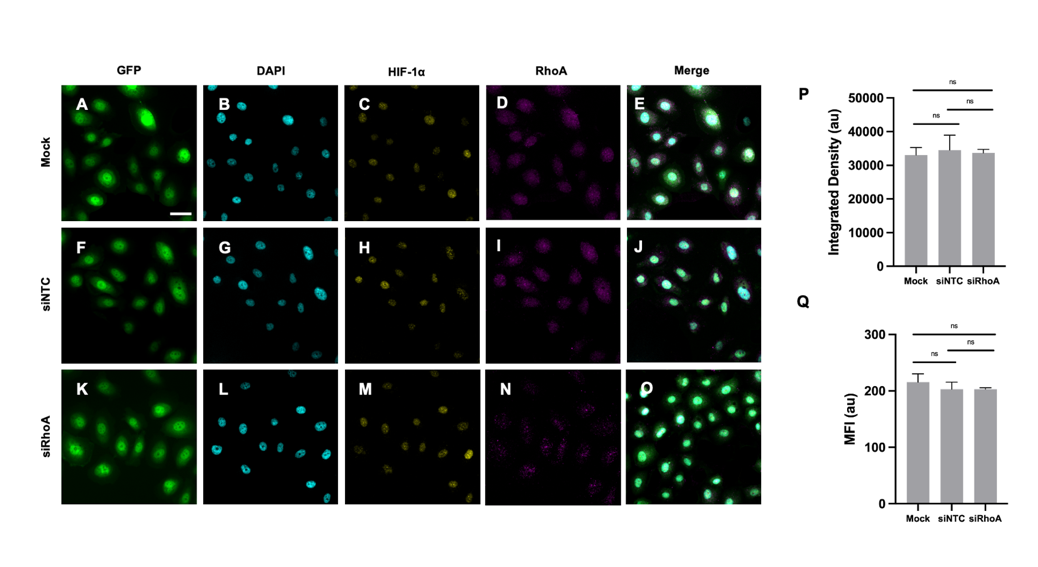

**Supplementary Fig. S6. HIF-1α expression is independent of RhoA in prostate cancer cells under normoxic conditions (A-O)** Immunofluorescence microscopy of PC3-GFP cells either mock transfected **(A-E),** transfected with non-targeting control siRNA **(**siNTC, **F-J),** or transfected with RhoA siRNA (siRhoA**, K-O)** under normoxic conditions. Panels show GFP **(A, F, K)**, DAPI **(B, G, L)**, HIF-1α **(C, H, M)**, and RhoA **(D, I, N).** Merged images of all channels per condition are displayed in **(E, J, O). (P–Q)** Quantification of HIF-1α expression by MFI **(P)** and Integrated Density **(Q)** in normoxic cells transfected with non-targeting control siRNA (siNTC), RhoA siRNA (siRhoA), or mock transfected (Mock). Data represent mean ± SEM. Significance determined as ns = not significant via One-way ANOVA with Tukey-Kramer post hoc test (see Materials and Methods). Scale bar in **A** represents 20 μm and applies to **A-O.**
